# Enhanced cell deconvolution of peripheral blood using DNA methylation for high-resolution immune profiling

**DOI:** 10.1101/2021.04.11.439377

**Authors:** Lucas A Salas, Ze Zhang, Devin C Koestler, Rondi A Butler, Helen M Hansen, Annette M Molinaro, John K Wiencke, Karl T Kelsey, Brock C Christensen

## Abstract

DNA methylation microarrays can be employed to interrogate cell-type composition in complex tissues. Here, we expand reference-based deconvolution of blood DNA methylation to include 12 leukocyte subtypes (neutrophils, eosinophils, basophils, monocytes, B cells, CD4+ and CD8+ naïve and memory cells, natural killer, and T regulatory cells). Including derived variables, our method provides up to 56 immune profile variables. The IDOL (IDentifying Optimal Libraries) algorithm was used to identify libraries for deconvolution of DNA methylation data both for current and retrospective platforms. The accuracy of deconvolution estimates obtained using our enhanced libraries was validated using artificial mixtures, and whole-blood DNA methylation with known cellular composition from flow cytometry. We applied our libraries to deconvolve cancer, aging, and autoimmune disease datasets. In conclusion, these libraries enable a detailed representation of immune-cell profiles in blood using only DNA and facilitate a standardized, thorough investigation of the immune system in human health and disease.

## Introduction

Advances in DNA methylation microarrays have allowed a greater understanding of how DNA methylation is affected by environmental exposures and altered in chronic diseases.^1,2^ Peripheral blood is one of the most common biological matrices for those investigations. Blood DNA methylation profiles include information from multiple cell lineages and, in some cases, cell states. Every cell lineage has unique DNA methylation patterning regulating cell-specific gene expression, and some methods leverage DNA methylation to understand underlying cell heterogeneity ^1–7^. The reference-based approach relies on the notion that the principal source of signal variability in a heterogeneous sample (such as blood) reflects the signals’ proportions in the different cell components ^8^. Constrained projection/quadratic programming (CP/QP) employs purified cell types as reference samples to generate a reference library of differentially methylated sites among cell types and yields highly accurate estimates of the underlying cell composition in mixed cell populations (e.g., peripheral blood)^9^. Previously established statistical deconvolution frameworks such as CP/QP, support vector machine (CIBERSORT), and robust partial regression (EpiDISH) have similar accuracy and precision in deconvolution estimates^10^. Marker selection methods for library creation use automatic procedures to discern library markers^11^ or iterative approaches for selecting sets of markers (IDOL) that maximize the accuracy of deconvolution estimates^12^. To enhance the utility of cell-type deconvolution, reference library improvements and expansions of existing libraries to include additional cell types are needed to broaden the scope of DNA methylation-based immune phenotyping.^13–16^

Continued methodological advancements are highly dependent on the quality and genome coverage of a reference library. In the original description of CP/QP for methylation-based deconvolution, Houseman et al. developed a library based on an early microarray platform, the Illumina HumanMethylation27k microarray ^9^. When the Illumina HumanMethylation450k technology was released, Jaffe et al. applied the Houseman method with the reference data developed by Reinius et al. ^11,17^. They accurately discriminated CD8 and CD4 T-cells, but NK and granulocytes (neutrophils and eosinophils) discrimination performance showed room for improvement^11,17^. Potentially limiting generalizability, the reference cell populations were purified solely from males of white Northern European origin ^17,18^. We recently developed a deconvolution library to better discriminate six major cell-types (CD4(+) T-cells, CD8(+) T-cells, NK, B-cells, monocytes, and neutrophils) using the Illumina HumanMethylationEPIC technology, hereafter named EPIC IDOL-6 ^19^. A distinct advantage of this library is the inclusion of more ethnically diverse male and female subjects.

Beyond the six major leukocyte cell types in peripheral blood, there have been further attempts to deconvolve memory and naïve cells and other granulocytes^13,14^. However, they have not been widely adopted or tested as some of the references are not publicly available or involved a combination of different technologies. Some algorithms include other rarer cell subpopulations [e.g., plasmablasts, exhausted CD8(+) T-cells] as linearly related scores, but they do not represent the sample’s cell-type proportions^15^. Several newborn umbilical cord blood-specific libraries also have been developed ^20–24^.

Here, we augment reference-based deconvolution of adult peripheral blood DNA methylation data to include memory and naïve cells from both cytotoxic and helper T-cells and B-cells and parse the granulocyte subtypes into neutrophils, eosinophils, and basophils (**Figure 1**). Our comprehensive library provides information across 12 different cell subtypes, resulting in 19 relative cell-type proportions and 19 cell-counts (derived data using the complete cell blood counts or flow cytometry). In addition, the library includes multiple cell-derived ratios and proportions such as the neutrophil to lymphocyte ratio or naïve to memory ratios (see **Figure 1** for 18 known examples), totaling more than 56 total immune profile variables. This library, hereafter named EPIC IDOL-Ext, will find wide application to the study of immune profiles in health and disease. Additionally, a second extended library using probes present on the legacy DNA HumanMethylation450k array, henceforth named 450k IDOL-Ext, was created for application to existing 450K datasets to expand cell-type representation.

**Figure 1.**
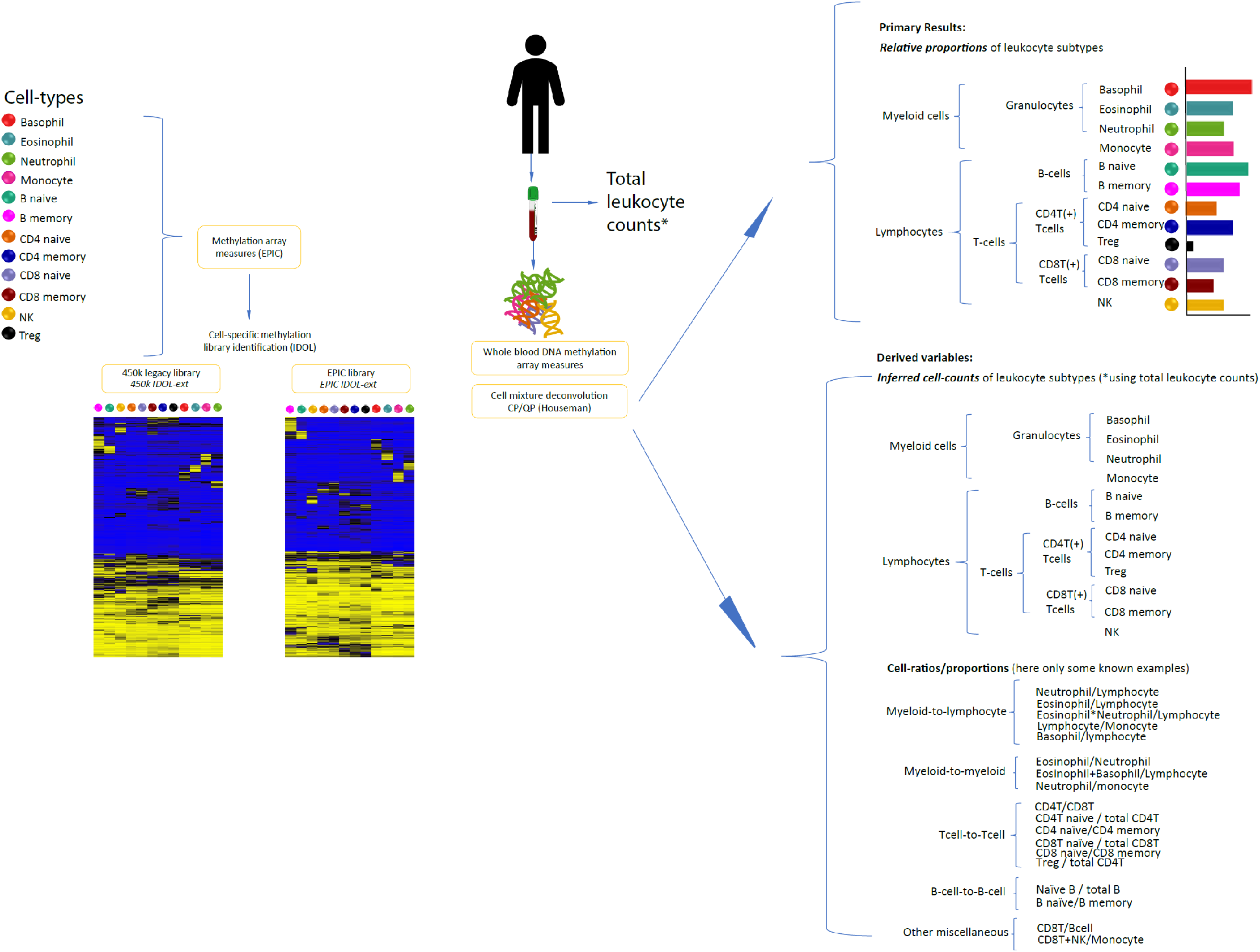
Summary of the process and expected results

## Results

To define a novel deconvolution library from 12 purified cell-types, we measured DNA methylation and performed rigorous quality assessment and control for all samples. The final reference data set included the following cell types: neutrophils (Neu, n=6), eosinophils (Eos, n=4), basophils (Bas, n=6), monocytes (Mono, n=5), B naïve cells (Bnv, n=4), B memory cells (Bmem, n=6), T-helper CD4+ naïve cells (CD4nv, n=5), T-helper CD4+ memory cells (CD4mem, n=4), T regulatory cells (Treg, n=3), T-cytotoxic CD8+ naïve cells (CD8nv, n=5), T-cytotoxic memory CD8+ cells (CD8mem, n=4), and natural killer cells (NK, n=4). The estimated purity of reference samples is based on commonly accepted CD marker definitions (**Supplementary Table 1**). The mean purity obtained from the flow cytometry confirmation step (after antibody-linked magnetic bead sorting) was 93% (range 85 to 99%), with the lowest purity observed for the CD8mem samples (85%). We first used minfi’s pickCompProbes function to select an automatic library and estimated cell-type proportions using this library with methylation data from purified cells. This library represents the expected average signal of some extreme hypo and hypermethylated markers per cell-type. Pure samples would approximate the average signal of the specific cell-type. However, when a sample is contaminated, the signal from other cell-type(s) will differ from the average and indicate the potential contaminant’s proportion. This procedure for reverse cell-type estimation is denoted as “DNA methylation purity.” We have used this technique previously to corroborate cell-identity and estimate potential residual cross-contamination during the flow cytometry procedures^24^. The mean DNA methylation purity for these samples was 97.6% (range: 85.7-100%), where a target cell type purity of 85% was required to include the sample in the dataset for library construction. Of the cell subtypes included, CD4nv had the highest estimated cell purity (median DNA methylation purity: 100%, interquartile range-IQR: 96.5-100%), and CD8mem cells had the lowest DNA methylation purity (median DNA methylation purity: 93%, IQR: 88.3-97.8%)). The remaining cell-types had median DNA methylation purity that ranged between 97.5 and 99.4%, IQR: 97.3-99.4%, **Supplementary Fig. 1**. Potential genetic sources of variability were assessed, including known SNPs tracing genetic ancestry (**Supplementary Fig. 2**). We excluded probes potentially tracking to polymorphisms, cross-reactive areas, or CpHs, probes tracking to sex chromosomes, and those whose signal intensities were equal or below to the background probes (see Online Methods for details). After filtering, 675,992 high-quality probes were retained for analysis and deconvolution library construction. The first 20 principal components showed that the main sources of variability corresponded to the cell-type identity and the slide beadchip, and not sex, age, or other phenotype variables (**Supplementary Fig. 3**).

We next applied the IDOL algorithm to the 56 samples collected across all 12 cell types using the 675,992 high-quality CpGs to select the optimal library for accurate deconvolution of these 12 cell types. To test our library’s performance, we used artificial mixtures (n=12) of DNA from purified cell types representing varying proportions of the 12 cell-types in the IDOL analysis and measured DNA methylation in these samples (**Figure 2a, Supplementary Table 2**). Using a discovery selection pool totaling 3,535 CpGs representing candidate markers of differentially methylated CpGs across the interrogated cell-types, we implemented the IDOL algorithm with 500 iterations to select a set of libraries ranging in size from 250 to 3000 CpGs, in increments of 50 CpGs (detailed in the methods). The coefficient of determination (R^2^) and the root mean square error (RMSE) were calculated based on a comparison of deconvolution estimates obtained using each library versus the known proportions of the 12 cell types in the artificial mixture samples (**Supplementary Table 3**). The optimal library, EPIC IDOL-Ext, consisted of 1,200 CpGs and was observed to have an average R^2^ of 1 across all cell types and the lowest average RMSE (0.226) among the interrogated library sizes. The training and testing datasets showed high R^2^ and low RMSE across the different interrogated cell-types (**Figure 2b**). Probes in the EPIC IDOL-Ext library tracked mostly to open sea regions of low CpG density (76%), followed by north and south CpG island shore regions (14%) (see **Table 1**). The vast majority of probes were tracked to DNAse hypersensitivity sites-DHS (73%). Notably, the library probes were also highly distinct as only 4% overlapped with the EPIC IDOL-6^19^.

**Table 1.** Characteristics of the context of the different blood cell-type deconvolution libraries

|  | EPIC IDOL-Ext<br>n=1200<br>N (%) | 450k IDOL-Ext<br>n=1500<br>N (%) | EPIC IDOL-6<br>n=450<br>N (%) | pickCompProbes<br>n=1200<br>N (%) |
| --- | --- | --- | --- | --- |
| Genomic context |  |  |  |  |
| CpG Island | 50 (4) | 122 (8) | 10 (2) | 62 (5) |
| North and South Shores | 189 (16) | 388 (26) | 63 (14) | 146 (12) |
| North and South Shelves | 95 (8) | 213 (14) | 35 (8) | 90 (8) |
| Open Sea | 866 (72) | 777 (52) | 342 (76) | 902 (75) |
| Functional context* |  |  |  |  |
| TSS1500 | 104 (9) | 192 (13) | 38 (8) | 83 (7) |
| TSS200 | 32 (3) | 81 (5) | 11 (2) | 40 (3) |
| 5'UTR | 128 (11) | 169 (11) | 46 (10) | 118 (10) |
| 1<sup>st</sup> Exon | 15 (1) | 31 (2) | 7 (2) | 28 (2) |
| Exon Band | 14 (1) | 0 (0) | 1 (0) | 5 (0) |
| Body | 569 (47) | 650 (43) | 209 (46) | 589 (49) |
| 3'UTR | 35 (3) | 77 (5) | 9 (2) | 31 (3) |
| Intergenic | 303 (25) | 300 (20) | 129 (29) | 306 (26) |
| Enhancers | 139 (12) | 76 (5) | 70 (16) | 194 (16) |
| DHS | 856 (71) | 1001 (67) | 328 (73) | 868 (72) |
| Open chromatin | 135 (11) | 174 (12) | 43 (10) | 113 (9) |
| TFBS | 166 (14) | 193 (13) | 59 (13) | 147 (12) |
| Contained in 450K | 459 (38) | 1500 (100) | 149 (33) | 517 (43) |
| Total overlap vs. Ref. |  |  |  |  |
| vs. EPIC IDOL-Ext | Ref | 330 (22) | 43 (10) | 147 (12) |
| vs. 450k IDOL-Ext | 330 (28) | Ref | 27 (6) | 89 (7) |
| vs. EPIC IDOL-6 | 43 (4) | 27 (2) | Ref | 57 (5) |
| vs pickCompProbes | 147 (12) | 89 (6) | 57 (13) | Ref |

\*Only first transcript reported
Acronyms: TSS1500 and TSS200 1500 and 200 base pairs upstream of the transcriptional starting site, 5'UTR and 3'UTR 5' and 3' untranslated region, DHS DNase Hypersensitive site, TFBS transcription factor binding site (as reported by the EPIC annotation), Enhancers (according to Phantom 5 in the EPIC annotation file), Ref reference for comparison in the column.

**Figure 2.**
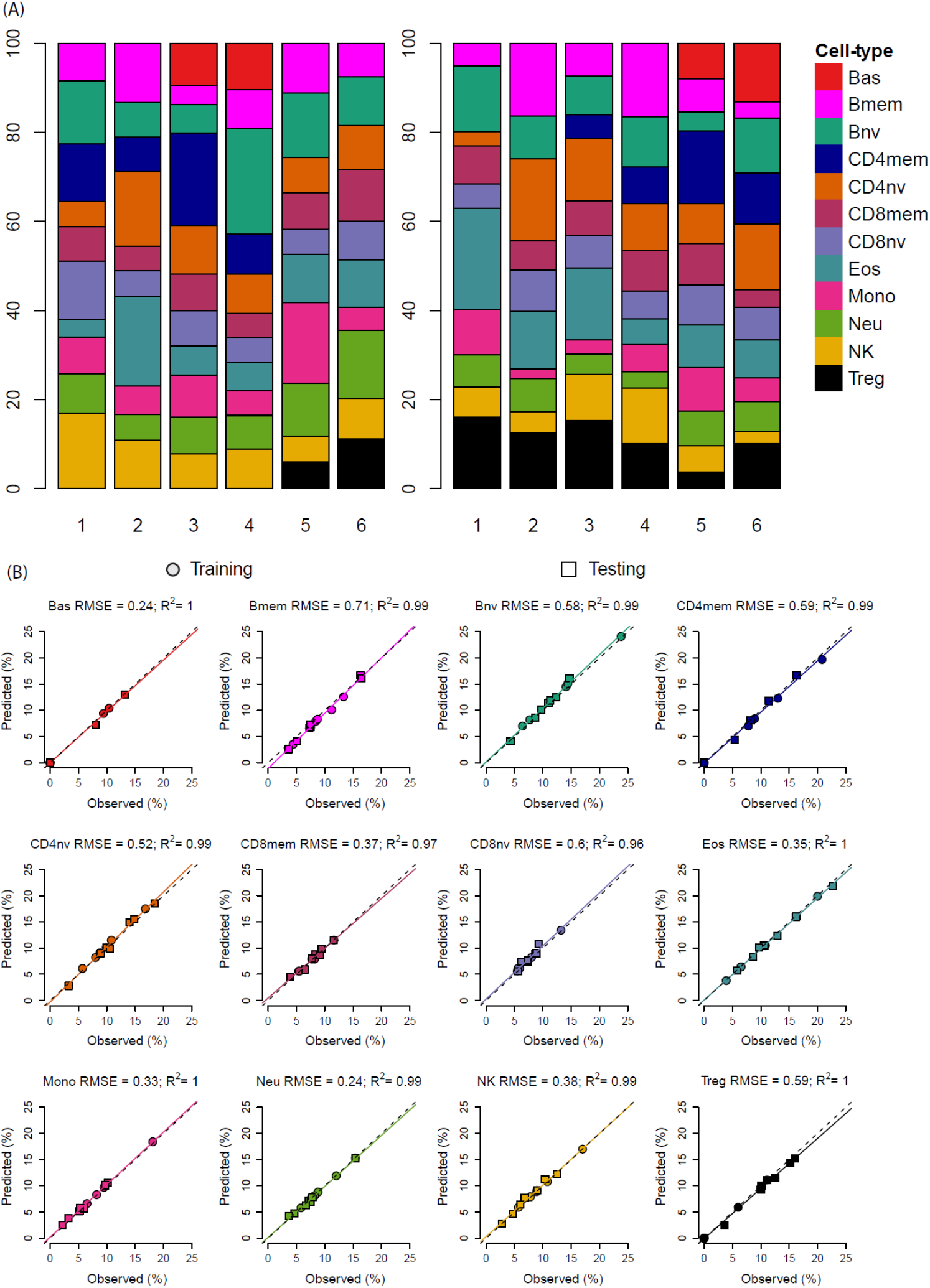
Comparison of estimate cell proportions using constrained projection/quadratic programming (CP/QP) versus the reconstructed (true) DNA fraction in the artificial DNA mixtures using the EPIC IDOL-Ext method. (a) Cell-specific DNA proportions per sample included in the training and testing sets. (b) R2 and RMSE using the EPIC IDOL-Ext method in both sets

As only 459 (38%) of the 1,200 probes in the EPIC IDOL-Ext library are common to both the EPIC and 450k array platforms, we developed a library for the legacy IlluminaHumanMethylation450k array platform, repeating the above-described optimization process after constraining the selection pool for the candidate list of CpGs to those only present on the 450k array. The resulting library, 450k IDOL-Ext, contains 1,500 probes; the library details are included in **Table 1**. Consistent with a different distribution of genomic contexts for probes on the 450k array, the 1,500 probe library had a lower proportion of open sea (52%) and enhancer-related probes (5%). In total, 330 probes were shared between the 450k IDOL-Ext and EPIC IDOL-Ext libraries.

Finally, we compared the EPIC IDOL-Ext, and 450k IDOL-Ext versus the pickCompProbes EPIC library obtained using functions in the minfi Bioconductor package^11^. The pickCompProbes automatic selection method builds the library picking the top 50 most hyper- and hypomethylated CpGs per cell-type, totaling 1,200 probes (same size as our EPIC IDOL-Ext library), summarized in **Table 1**. The overlap of probes from the pickCompProbes library with the EPIC IDOL-Ext library was only 147 probes (12%), though the probes’ genomic context distribution was similar between libraries. When analyzing the cell-type by cell-type estimation performance across all three libraries, there was consistency in monocytes, neutrophils, and NK cells. However, more variability was observed for estimates obtained from the library derived by pickCompProbes. When assessing the deconvolution accuracy of the pickCompProbes library, the RMSE was severely biased for eosinophils and T-cell subtypes. Specifically, CD4 and CD8 naïve T-cell distributions were biased with an underrepresentation of CD8nv (RMSE: 5.81%) and the overestimation of CD4nv (RMSE: 9.25%). The pickCompProbes library also had unreliable results for CD4mem vs. Treg compartments and Eos, **Supplementary Fig. 4**. In contrast, both the EPIC IDOL-Ext and 450k IDOL-Ext libraries were highly accurate in the training and testing datasets, **Figure 2** and **Supplementary Fig. 4**. The heatmaps summarizing the markers in the three libraries are shown in **Figure 3**. The complete set of markers information is available as **Supplementary Tables 4** (EPIC IDOL-Ext), **5** (450k IDOL-Ext), **and 6** (pickCompProbes).

**Figure 3.**
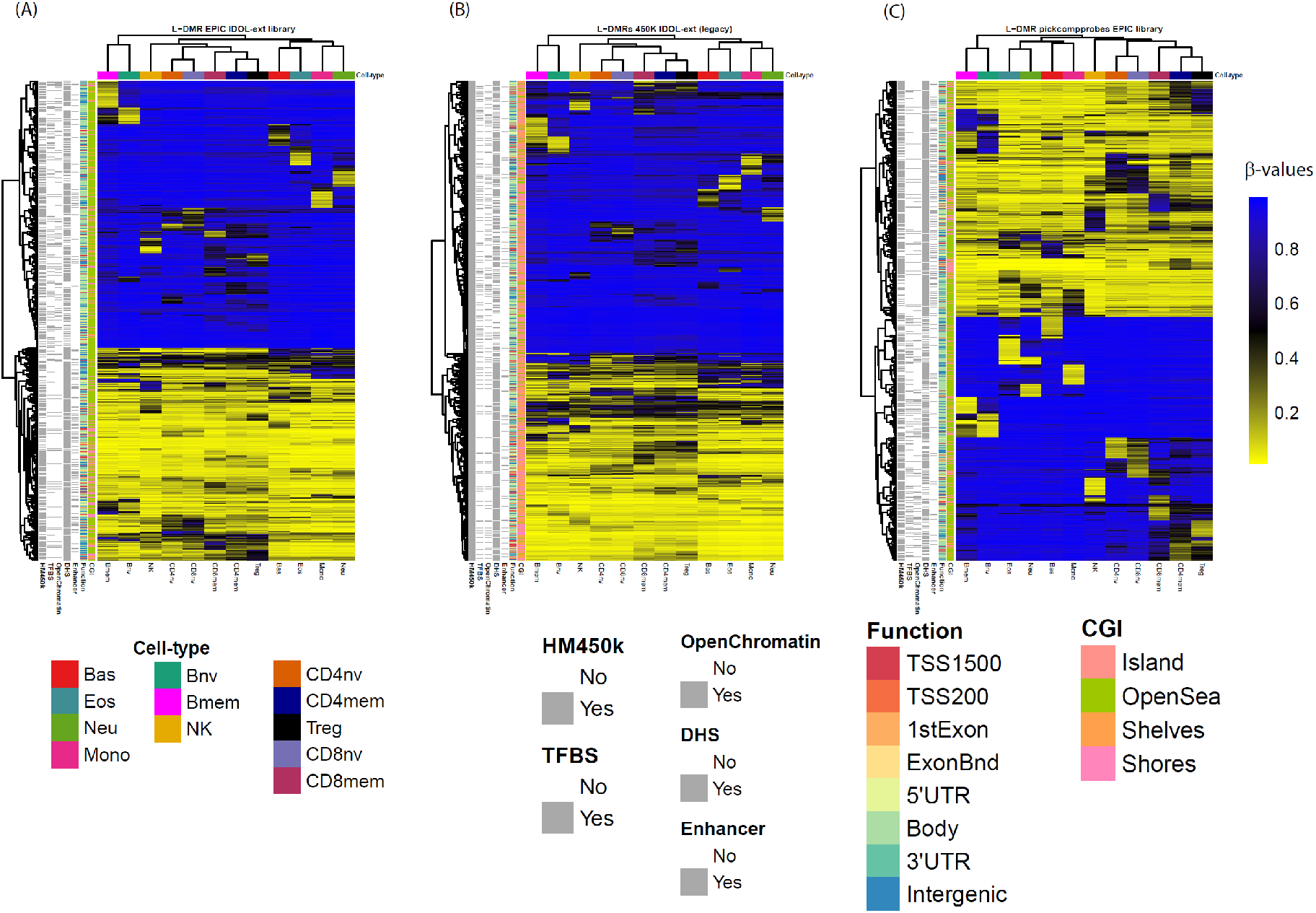
Comparison of the selected CpG among the EPIC IDOL-Ext, 450k IDOL-Ext (legacy), and pickComProbes. (a) DNA methylation of the 1200 CpG probe EPIC IDOL-Ext library with average methylation for cell type samples shown in columns as labeled and tracking bars for CpG probes in rows including present on 450k array, tracking to transcription factor binding site (TFBS), DNA hypersensitivity site (DHS), enhancer region from Phantom5 annotation, and genomic context relative to gene (Function) and CpG island (CGI). (b) DNA methylation of the 1500 CpG probe 450k IDOL-Ext (legacy) library and (c) DNA methylation of the 1200 CpG pickComProbes library. Note: the average DNA methylation levels (here as beta values) are represented per each of the 12 cell-types. Information about CpG island gene context, Phantom5 enhancer information, DNase Hypersensitivity sites (DHS), Open chromatin, and annotated transcription factor binding sites (TFBS) from ENCODE and the Illumina annotation file are summarized using the row ribbons.

The EPIC IDOL-Ext library was validated using samples with blood cell counts from flow cytometry (FCM) and by using an independent set of artificial mixtures from the Gene Expression Omnibus (GEO) (**Figure 4**). In healthy adult samples (**Figure 4a**, GSE110530), we observed a strong correlation between the deconvolution estimates and FCM measurements, with a maximum root mean square error-RMSE of 3.29% for CD8T cells. Using independent artificial mixtures (**Figure 4b**, GSE110554), the correlation was close to 1, and the maximum RMSE was 1.67 for the neutrophils (granulocytes). The 450k IDOL-Ext library was also validated using the GSE77797 with FCM (**Figure 4c**) and artificial mixture information (**Figure 4d**).

**Figure 4.**
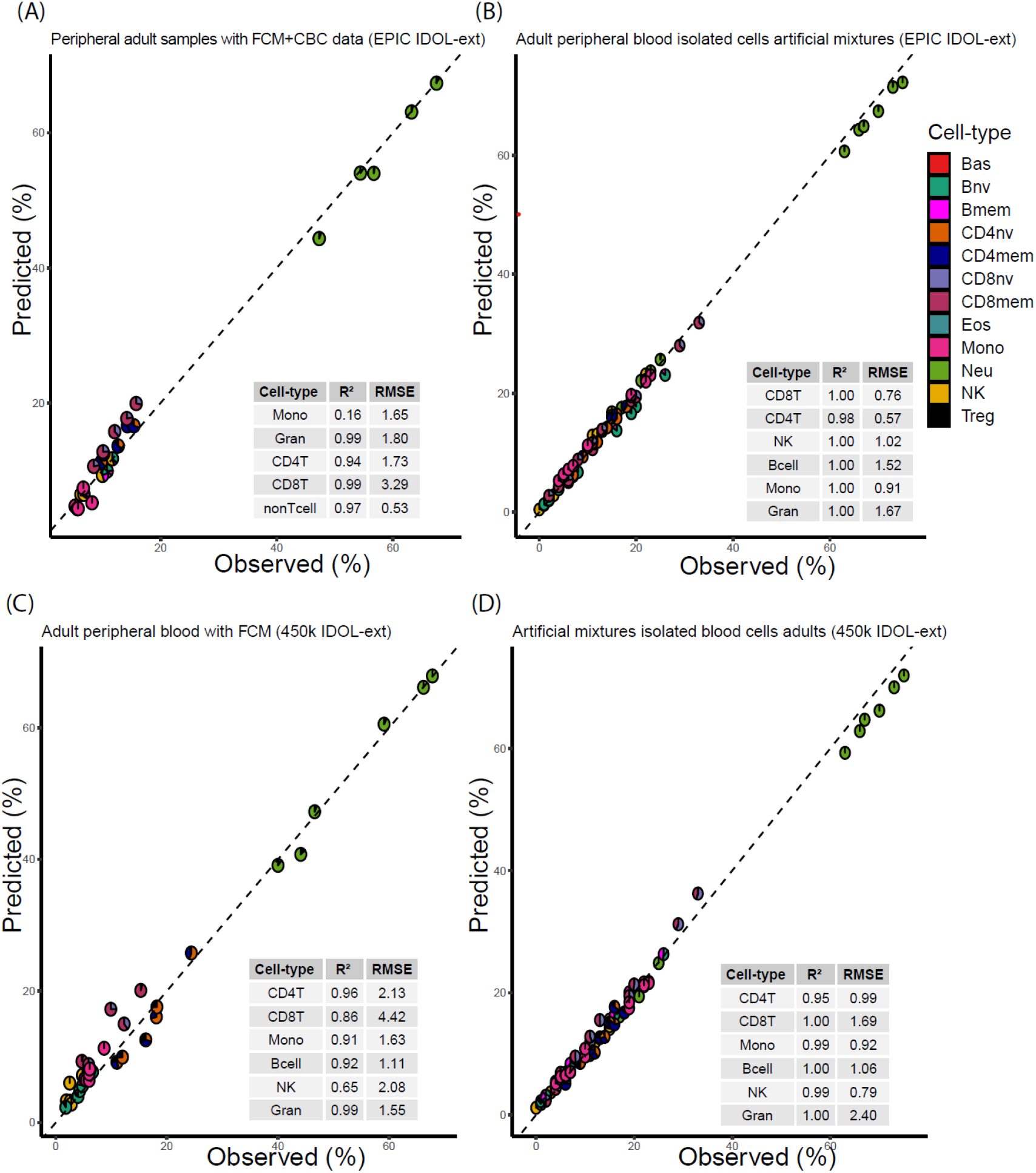
Validation of the library using flow cytometry-FCM and independent artificial mixtures for adult EPIC IDOL-Ext (a and b) and 450k IDOL-Ext (c and d) Notes: The area of each pieplot corresponds to the estimated proportion of the cell-types within each group. Gran (Granulocytes) corresponds to the sum of Neu-neutrophils, Eos-eosinophils, and Bas-basophils. CD4T corresponds to the sum of CD4+ T cells naïve, memory and Treg. CD8T corresponds to the sum of CD8+ T cells naïve and memory. Bcell to the sum of the naïve and memory. No basophils were detected in the samples illustrated in (a). Additionally in (a), “nonTcell” (Bnv, Bmem and NK) were not experimentally measured.

As a proof-of-concept, we explored how the estimates could help reconstruct cell counts using FCM information (**Supplementary Fig. 5**). Additional validation sets were analyzed, including glioma patient blood and the cytometric information for the Reinius dataset^17^. A group of 72 glioma patients was used to evaluate the information from T-cell subsets. For this, cytometric information was obtained using a two-stage characterization of T cell subtypes. In the first stage, T-cells were measured and separated into CD4+ and CD8+. Next, using a second tube, samples were characterized as naïve or memory. Although the samples were measured in the same subjects, the proportions were estimated based on two independent measurements and were mathematically derived based on both experiments’ average counts. We observed the second-largest difference for the CD8mem with an RSME of 5.44% and the largest difference with the unmeasured remainder cells (RMSE of 12.48%) (**Supplementary Fig. 6a**). The 450k IDOL-Ext was also validated using the GSE35069 datasets (Reinius) with FCM information (**Supplementary Fig. 6b**). We observed considerable variation (∼10%) for the granulocytes, monocytes, and CD4T cells in this dataset ^17^. Differences were more prominent for the PBMC samples’ observed values and in two of the six WBC samples, particularly for the granulocytes reported.

Finally, we explored whether these libraries could be applied for the deconvolution of umbilical cord blood samples. We used the GSE68456 dataset (450k) with known FCM information for seven cell-types, including the erythroblasts (nucleated red blood cells). As this subtype was not present in our library, the fraction was added to the myeloid granulocyte components for comparison. This fraction showed the largest variability in this dataset with an RMSE of 15.57% (**Supplementary Fig. 7a**). In contrast, when using umbilical cord blood artificial mixtures excluding the nucleated red blood cells, the largest RMSE was observed for granulocytes being less than 4% (**Supplementary Fig. 7b)**.

The different libraries’ components were examined for how the included genes were related to specific cell-types and immune pathways. First, we used the molecular signatures database (MSigDB), curated by the Broad Institute, for gene set enrichment analyses v. 7.2. Most of the pathways were immune-related and are summarized in **Supplementary Table 7**. Some specific genes showing differential hypomethylation for specific cell-types are outlined in **Figure 5**.

**Figure 5.**
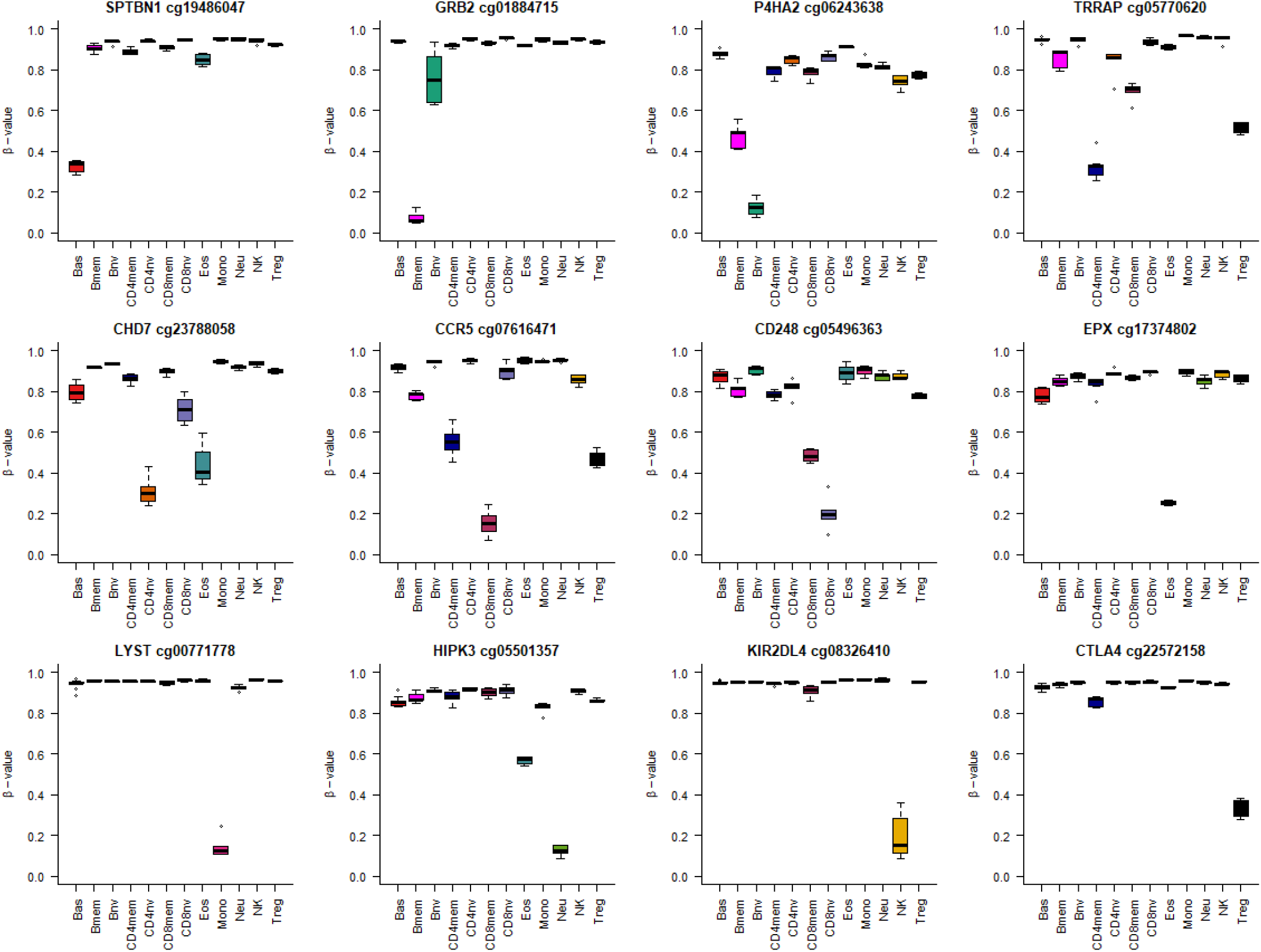
Examples of differentially methylated sites demarcating the different cell lineages

We next applied the EPIC IDOL-Ext library and the 450k IDOL-Ext libraries to several publicly available datasets from GEO and ArrayExpress to identify potential variation in immune cell proportions in multiple sclerosis, rheumatoid arthritis, breast cancer patients, and COVID-19 infection (**Supplementary Table 8**). In multiple sclerosis patients, we observed significantly increased basophil and naïve B cell proportions in cases (n=13) compared to controls (n=14) (Wilcoxon rank-sum *P <0*.*01*, **Supplementary Fig. 8)**. In rheumatoid arthritis, significant increases of neutrophil and regulatory T cell proportions and decreases of memory B cell, memory CD4 cell, naïve CD4 cell, memory CD8 cell, naïve CD8 cell, eosinophil, monocyte, and NK cell proportions were observed in case blood samples (n=354) compared to control blood samples (n=355) (Wilcoxon rank-sum *P <0*.*01*, **Supplementary Fig. 9)**. Predicted immune cell proportions in breast cancer patients before and after receiving chemotherapy or a combination of chemo/radiation therapy were explored. Patients receiving radiation therapy only (n=74) showed a significant relative increase of the neutrophil proportion and a mirror decrease of several lymphoid lineages (memory B cell, naïve B cell, naïve CD4 cell, and NK) after treatment (Wilcoxon rank-sum *P <0*.*01*, **Supplementary Fig. 10a)**. Patients receiving radiation therapy and chemotherapy (n=70) exhibited significant increases in eosinophil, monocyte, and regulatory T cell proportions and decreases of memory B cells after treatment (Wilcoxon rank-sum *P <0*.*01*, **Supplementary Fig. 10b)**. Finally, we evaluated the changes in immune cell proportions in six COVID-19 patients with and without remission versus six healthy controls in **Supplementary Fig. 11**. Because of the limited sample size, no statistically significant differences were observed, but, as expected, the median of neutrophils was higher in patients versus controls. In contrast, all the median lymphocyte subpopulations were lower in the infected patient than those who recovered from the disease. The median monocytes were lower in those that remitted.

We investigated blood methylation data associated with subject-to-subject variation in non-pathological conditions using data from monozygotic versus dizygotic twins and subjects at different ages (**Supplementary Table 8**). To characterize immune cell variation between twins, predicted immune cell proportions in monozygotic twins (n=852) and dizygotic twins (n=612) were estimated. Significant differences in all the immune cells were observed in monozygotic twins and dizygotic twins (Paired t-test *P<0*.*01*). Larger differences between dizygotic twins compared to monozygotic twins were observed in memory B cell, naïve B cell, memory CD4 cell, naïve CD4 cell, memory CD8 cell, naïve CD8 cell, eosinophil, monocyte, and NK (Wilcoxon rank-sum *P <0*.*01*, **Supplementary Fig. 12)**. Next, the impacts of aging on predicted immune cell proportions in blood samples from newborn to nonagenarian (n=2504) were investigated. Several subpopulation cell ratios across different ages were calculated **(Supplementary Fig. 13)**. The complete granular data and trajectories of the different cell subpopulations are represented in **Supplementary Fig. 14**. Longitudinal changes of predicted immune cell proportions within five years after birth in human blood leukocytes from 10 healthy girls are shown in **Supplementary Fig. 15**.

## Discussion

We established compact and reliable libraries to deconvolve the proportions of 12 different cell-types in peripheral blood, including closely related cell-types such as T-cell subtypes and various types of granulocytes. Importantly, our libraries’ derived variables offer a detailed immune profile from peripheral blood, providing more than 56 cell-type and ratio components. Our libraries are designed both for current and legacy DNA methylation measurement platforms that together include publicly available data from >100,000 samples. Our libraries have high accuracy and minimal bias when compared to flow cytometry gold standard data. The new libraries will be of importance where specific memory and naïve cell compartments are interrogated for disease, pathogenesis, or exposures.

We consistently observed biologically relevant genes and groups of genes in related pathways for the different cell-types, such as the *SPTBN1*, which is more expressed in basophils ^25^; *GRB2*, involved in Bmem formation ^26,27^; *P4HA2*, related to immature B cells; *CHD7*, associated with thymic function ^28^; *CCR5*, related to CD8mem recruitment and trafficking ^29^; CD248 a CD8 regulator and marker of naïve states ^30^; *EPX*, the eosinophil peroxidase marker of eosinophils ^31^; *LYST* a lysosome maturation in monocytes ^32^; or *KIR2DL4* a natural killer cell receptor ^33^; and the *CTLA4* an essential marker in Treg tumor suppression.^34^

Our EPIC IDOL-Ext and 450k IDOL-Ext libraries provide enhanced features for researchers using DNA methylation data derived from blood. When the research question requires finer resolution, the newly extended libraries provide that extra information, with a minimal trade-off in precision compared with the earlier EPIC IDOL-6 library. Previous attempts to generate extended libraries for blood deconvolution encountered several problems. The discrimination of scarcer cells has been problematic, leading to inconsistent estimates, even when candidate references were available. Some markers delineating particular cell-types are dependent on sex chromosome dosage (e.g., *FOXP3* located in the X chromosome for Tregs). To avoid that potential problem here, we only used autosomal genes. Distinguishing cell-types hierarchically close (i.e., CD4mem and Tregs) is challenging as the pool of potential specific markers is limited. These markers may even show some dependency when transitioning from one state to another. For example, *CD248* is mostly unmethylated in CD8nv, but some cells in the CD8mem compartment are not completely methylated at *CD248*, while it is completely methylated and repressed in other cell-types. In such case, the use of multiple markers attenuates the effect of using single markers to define cell-identity. Finally, when using libraries derived from automatic selection, the statistical process may favor markers that do not necessarily reflect biologically relevant genes. For example, in the extended libraries, four (EPIC) and five (450k) *EPX* (eosinophil peroxidase) probes were selected, in contrast to none by the minfi pickCompProbes procedure.

The inclusion of six additional cell-types and, in particular, of the naïve compartments is a major step towards a universal deconvolution library applied to cord or adult blood data. Particularly intriguing for umbilical cord blood, nucleated red blood cells interfered with more precise estimations than FCM information, yet artificial mixtures showed the library capturing leukocyte components accurately. As expected, most (if not all) of the B and T-cells were naïve cells as memory populations are derived after antigen exposures, most commonly after birth.

Another advance of this library is the most accurate eosinophil predictions compared to previous libraries and now the inclusion of basophil estimations. These cells are involved in multiple pathologies, including allergies and asthma, and through these libraries can now be estimated. As with previous libraries, some residual confounding from unaccounted cell-types may be observed. In particular, we observed inflation in several datasets of the basophil estimation. Thus careful interpretation based on those estimates alone is prudent. Based on single-cell murine data, one hypothesis for this estimate inflation is common progenitors’ relation with triple differentiation to basophil/mast cells and erythroid cells ^35,36^. According to those potential common markers, it is possible the basophil signal could interfere with the signal derived from circulating immature nucleated erythroid cells. At least in umbilical cord blood samples, the signal obtained from basophils is highly correlated with the predicted fraction of nucleated red blood cells. As such, a careful assessment of any high basophil signal is warranted, as it could be driven by interference with nucleated red blood cells in newborns and potentially some pathological or adaptative conditions at other ages.

In summary, the new reference libraries greatly enhance the detail of immune cell profiling with DNA methylation. These enhanced libraries can be applied directly for immune profiling and adjustment of cell-type proportions in EWAS. Future work includes validating the use of this library in methylation data from children and umbilical cord blood. The increased detail available in describing leukocyte subtypes using the extended libraries will be crucial for controlling aging-related effects; age-associated changes in specific subpopulations of memory T and B lymphocytes unaccounted for previously established libraries. Indeed, the IDOL-Ext libraries’ application developed here greatly enhances the detail of immune characterization to existing data of immune profiles and in future studies using DNA methylation data derived from human blood samples.

## Methods

This work extended the available six-cell reference library using twelve cell-subtypes for deconvolution of blood cell proportions using the EPIC array (as well as a legacy library for the 450k platform). Using magnetic-sorted and flow-sorted confirmed neutrophils, eosinophils, basophils, B cells (naïve and memory), monocytes, NK cells, CD4+ T cells (naïve, memory and T regulatory cells), and CD8+ T cells (naïve and memory), we measured DNA methylation with the 850K/EPIC DNA methylation array. We applied the IDOL method to identify optimal Leukocyte -Differentially Methylated Regions (L-DMR) libraries. We then compared the performance of cell estimates obtained applying our previously developed library with six cell-types available in FlowSorted. Blood.EPIC and optimized with the L-DMR IDOL library to the older 450 K array in artificial blood mixtures with predefined cell proportions. Four MACS-isolated and FACS-verified purity cell sub-types from the myeloid lineage [neutrophils (Neu), eosinophils (Eos), basophils (Bas), and monocytes (Mono)], plus eight MACS-isolated and FACS-verified purity cell sub-types from the lymphoid lineage (B lymphocytes naïve (Bnv), B lymphocytes memory (Bmem), T helper lymphocytes naïve (CD4nv), T helper lymphocytes memory (CD4mem), T regulatory cells (Treg), T cytotoxic lymphocytes naïve (CD8nv), T cytotoxic lymphocytes memory (CD8mem), and natural killer lymphocytes (NK)) were purchased from AllCells^®^ corporation (Alameda, CA, USA) and STEMCELL technologies (Vancouver, BC, Canada). Cells were isolated from 41 males and 15 females, all anonymous healthy donors. The donors had a mean age of 32.2 years (range 19–58 years) and an average weight of 85.1 kg (range 57–136 Kg) and were negative for HIV, HBV, and HBC. Women were not pregnant at the time of sample collection, and samples were collected from donors with no history of heart, lung, kidney disease, asthma, blood disorders, autoimmune disorders, cancer, or diabetes. All donors provided written informed consent before donation. The data discussed in this publication have been deposited in NCBI’s Gene Expression Omnibus (Salas et al., 2021) and are accessible through GEO Series accession number GSE167998 (https://www.ncbi.nlm.nih.gov/geo/query/acc.cgi?acc=GSE167998). Isolation protocols are available through the commercial websites of AllCells and STEMCELL technologies. In brief, cells were selected using immunomagnetic labeling through the vendors’ specific protocols (see, **Supplementary Table 1** for details). Recovered cells were confirmed using flow-sorting. Twelve artificial mixtures were reconstructed using DNA from the specific cell samples. The reconstruction mixtures were determined by randomly generating proportions from a twelve-component Dirichlet distribution. The reconstructed samples used mixtures of purified leukocyte subtype DNA on relatively equivalent proportions across the twelve leukocyte subtypes. Each mixture contained 1.2 μg of total DNA was estimated using the proportions included in **Supplementary Table 2**. The DNA from the cell-sorted samples and those of the artificial mixtures were randomized in the slide slots of the micro-array. Sample DNA (1μg) was bisulfite converted and processed according to the Illumina protocols at the Vincent J. Coates Genomics Sequencing Laboratory at UC Berkeley, Avera, or Diagenode. The EPIC methylation arrays’ raw idat files were pre-processed using minfi, EnMIX, and SeSaMe for quality control using R v.4.0.2 ^37–39^. To assess data quality, we used an out-of-band detection P-value of 0.05, three standard deviations of the mean bisulfite conversion control probe fluorescence signal intensity, and a minimum of three beads per probe. To ascertain the highest purity of the samples included in our library, in addition to the information obtained through the FCM confirmation, we projected back the proportions of the cells in our library. This is to observe how homogeneous they were using Jaffe’s procedure^11^. The corresponding cell-type proportion was retrieved and designated as “DNA methylation purity.” We only included in this library samples with DNA methylation purity levels higher than 85% (range 85.7% to 100%). Next, we applied a stringent out-of-band p-detection value (pOOBHA) > 0.05 and set those that could not be distinguished from the background probes as missing values. Then we selected only those probes with complete information for all the samples in the library. No imputation was performed in this context as the signals could be dependent on the specific cell-type. Further, as we only observed a minimal variation in probes tracing to non-CpG or CpH probes, all the CpH beta values were under 0.08, they were filtered. Finally, as both females and male samples were present, we discarded probes tracking the X and Y chromosomes. According to Zhou et al., those that showed known polymorphisms or cross-reactivity were also excluded.^40^. Our set for library discovery included 675,992 complete high-quality probes.The EPIC IDOL-Ext L-DMR library is available in GitHub (https://github.com/immunomethylomics/FlowSorted.BloodExtended.EPIC), and will be available through Bioconductor with the name FlowSorted.BloodExtended.EPIC. The extended blood deconvolution can be performed using the FlowSorted.Blood.EPIC Bioconductor library. The package contains an RGChannelSet R object processed using SeSaMe in which probes showing channel switching were corrected and SNPs derived from Infinium Type I probes were added using the total signal intensities. The object is unfiltered and contains 56 samples and the 12 artificial mixtures information on 1,008,711 probes corresponding to 866,091 CpGs using the latest annotation released by Illumina (MethylationEPIC_v-1-0_B5). The reader needs to note that the cells were purified using an immunomagnetic procedure; the name “FlowSorted” is kept for historical reasons and downstream integration with previous minfi pipelines and similar algorithms.

### IDOL algorithm

For a complete description of the IDOL algorithm, please refer to the original application in Koestler et al. ^12^. In brief, the IDOL algorithm utilizes a training dataset (ground truth) consisting of samples with DNA methylation data in which the measured fraction of each of the underlying cell types is known (here corresponding to artificial mixtures with pre-specified proportions) as a means to identify a set of probes confirming an optimal reference library for cell mixture deconvolution. A series of t-tests compared the mean CpG-specific methylation between each leukocyte cell-type vs. the mean methylation across all the other cell-types identified the probes discriminating CpGs (e.g., leukocyte differential methylated regions or L-DMRs) for each specific cell-type of the 12 included in this application. CpGs were then rank-ordered using their t-statistics, and the *L*/2 CpGs with the largest and smallest t-statistic for each *K* cell type were identified and pooled. The tuning parameter *L* was set to *L*= 150 in our application, consistent with Koestler et al.^12^. A discovery L-DMR library containing the total *L*K* unique L-DMRs for each cell type forms the IDOL algorithm search space. L-DMR subsets of size <*L*K* are sequentially selected and examined for their prediction accuracy in deconvolving the training dataset samples. The user needs to preselect the library size to balance the accuracy and precision of cell composition estimates. For the current application of IDOL presented here, we considered libraries ranging from 250 to 3000 CpGs. We initially set increments of 50 CpGs until a size of 1100; then we increased by 100 CpGs until a size of 2000 and tested libraries of 2500 and 3000 CpGs to corroborate the elbow in the error distribution. In the first iteration of the IDOL algorithm, all *L*K* CpGs constituting the candidate library have an equal probability of being selected to be included in the L-DMR library. We applied the constrained projection/quadratic programming approach ^9^ to obtain the cell composition estimates for each sample in the training dataset. These predictions then allow us to calculate the R^2^ and RMSE (root mean square error) for each of the cell-types, contrasting the cell estimates versus the known proportion in each sample. Then one-by-one CpGs are removed from the randomly selected DMR library, followed by computation of R^2^ and RMSE based on cell-composition estimates obtained using the new library. Through this, we can assess the contribution of each CpG in the library in terms of its impact on the accuracy of cell composition estimates and, then the algorithm modifies the probability of each CpG being selected in subsequent IDOL iterations. The process is repeated at each of the 500 iterations, with the algorithm eventually converging on an “optimal” library for deconvolution (showing the lowest error and highest precision).

### Enrichment analysis

Gene set enrichment analyses used missMethyl to control for multiple probes bias and the Molecular signatures database version 7.2. ^41,42^

### Validation sets

We used five datasets for validation: a set of samples obtained from a longitudinal analysis (GSE110530) with six observations. A set of independent artificial mixtures (GSE110554) derived from six cell-types. A set of samples from 72 glioma patients in different treatment stages, including FCM information for T cells CD4+ and CD8+ naïve and memory. A set of 20 umbilical cord blood (GSE68456) with FCM information. A set of 12 umbilical cord blood artificial mixtures including six cell-types. A set of samples using the 450k technology (GSE77797) with FCM and artificial mixture information derived from six cell-types. Finally, the Reinius et al. GSE35069 dataset using the FCM information for the whole blood cells, peripheral blood mononuclear cells, and granulocytes as reported in the original manuscript’s supplementary material.

### Potential applications

We identified 12 publicly available data sets from GEO that contained DNA methylation data on different health conditions and diseases (**Supplementary Table 8**). The application data sets included whole blood samples from 426 pairs monozygotic twins and 306 dizygotic twins, 2504 umbilical cord and peripheral blood samples from newborn to nonagenarian, 13 multiple sclerosis whole blood case samples and 14 controls, 354 rheumatoid arthritis peripheral blood leukocyte samples and 355 controls, 144 peripheral blood samples from breast cancer patients before and after receiving chemotherapy and radiation or isolated radiation therapy treatment, and six COVID-19 patients (with 18 samples) and six healthy controls. Illumina Infinium DNA methylation IDAT files were retrieved from GEO and ArrayExpress for application data sets. *SeSAMe* package from *Bioconductor* was used to process the data. Four thousand eight hundred and seventy-two samples in total were eventually contained in the application data sets. We estimated the proportion of immune cells in those data sets using the appropriate extended immune cell deconvolution library for the array (EPIC or 450k).

## Acknowledgments

L.A.S. was supported by CDMRP/Department of Defense (W81XWH-20-1-0778) and NIGMS (P20GM1044168299). D.C.K. was supported by NIH grants P30 CA168524, P20 GM130423, and P20GM103428. J.K.W. & A.M.M. were supported by NIH grants R01 CA207360 and P50 CA097257. K.T.K. was supported by a 2018 AACR-Johnson & Johnson Lung Cancer Innovation Science (18-90-52-MICH). B.C.C. was supported by NIH grants R01CA216265 and R01CA253976.

## Contributions

L.A.S., D.C.K., J.K.W., K.T.K., and B.C.C. conceived the project and designed the experiments. L.A.S., and Z.Z. performed the bioinformatic quality control and analyzed the data. H.M.H and R.B.B carried out the wet-lab work, including the spiking of the artificial mixtures used in this experiment (R.B.B.). D.C.K. and A.M.M. supported the statistical analysis. L.A.S wrote the manuscript with input from all the co-authors. The final version of the manuscript has been reviewed and approved by all the authors.

## Abbreviations

CP/QP: constrained projection/quadratic programming
FCM: flow cytometry
IDOL: improved cell deconvolution optimized library
L-DMR: leukocyte differentially methylated region
RMSE: root mean square error
Bas: Basophils
Eos: eosinophils
Neu: neutrophils
Bnv: B-cells naïve
Bmem: B-cells memory
CD4nv: helper CD4(+) T-cells naïve
CD4mem: helper CD4(+) T-cells memory
Treg: CD4(+) T regulatory cells
CD8nv: cytotoxic CD8(+) T-cells naïve
CD8mem: cytotoxic CD8(+) T-cells memory
Mono: monocytes
NK: natural killer cells.

