## Supplementary material for "Enhanced cell deconvolution of peripheral blood using DNA methylation for high-resolution immune profiling"

Supplementary Table 1 Cell-type definitions according to flow cytometry markers

| Group | Subgroup | Cell-type | Abbreviation | Selection | Markers | Source |
|---|---|---|---|---|---|---|
| Myeloid | Mononuclear cells | Monocytes (classical) | Mono | Negative | CD14(+) | PBMC |
| | Granulocytes | Basophils | Bas | Negative | IgE(+) CD123(+) | PMN rich fraction basophil labelling system |
| | | Eosinophils | Eos | Negative | CD15(+) CD16(-) | PMN rich fraction (HetaSep) |
| | | Neutrophils | Neu | Negative | NA | PMN rich fraction (HetaSep) |
| Lymphoid | T-cells | T regulatory cells | Treg | Negative | CD4(+) CD25(+) CD127dim/(-) | PBMC |
| | | T helper CD4+ naive cells | CD4nv | Negative | CD4(+) CD45RA(+) CD45RO(-) | PBMC |
| | | T helper CD4+ memory cells | CD4mem | Negative | CD4(+) CD45RA(-) CD45RO(+) | PBMC |
| | | T cytotoxic CD8+ naive cells | CD8nv | Negative | CD8(+) CD45RA(+) CCR7(+) CD45RO(-) CD56(-) CD57(-) | PBMC |
| | | T cytotoxic CD8+ memory cells (effector memory) | CD8mem | Positive | CD8(+) CD45RO(+) CD62(-) | PBMC |
| | B-cells | B naïve cells | Bnv | Negative | CD19(+) CD27(-) | PBMC |
| | | B memory cells | Bmem | Positive | CD19(+) CD27(+) | PBMC |
| | Natural killers | Natural killers | NK | Negative | CD56(+) | PBMC |

Supplementary Fig. 1. Cell purity estimated by flow sorting verification (panel A), and estimated DNA methylation purity (Panel B)

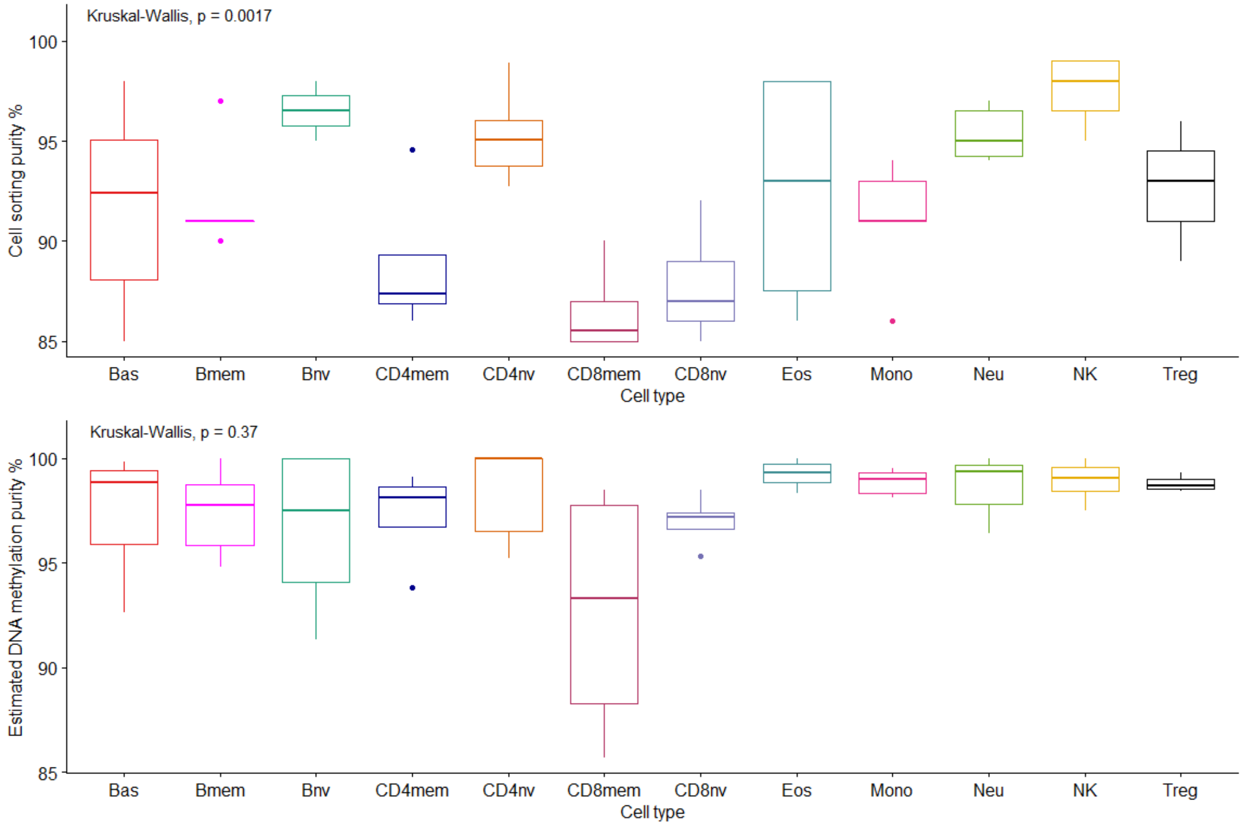

Supplementary Fig. 2. Known SNPs tracing to genetic ancestry markers distribution across the cell-types in the libraries

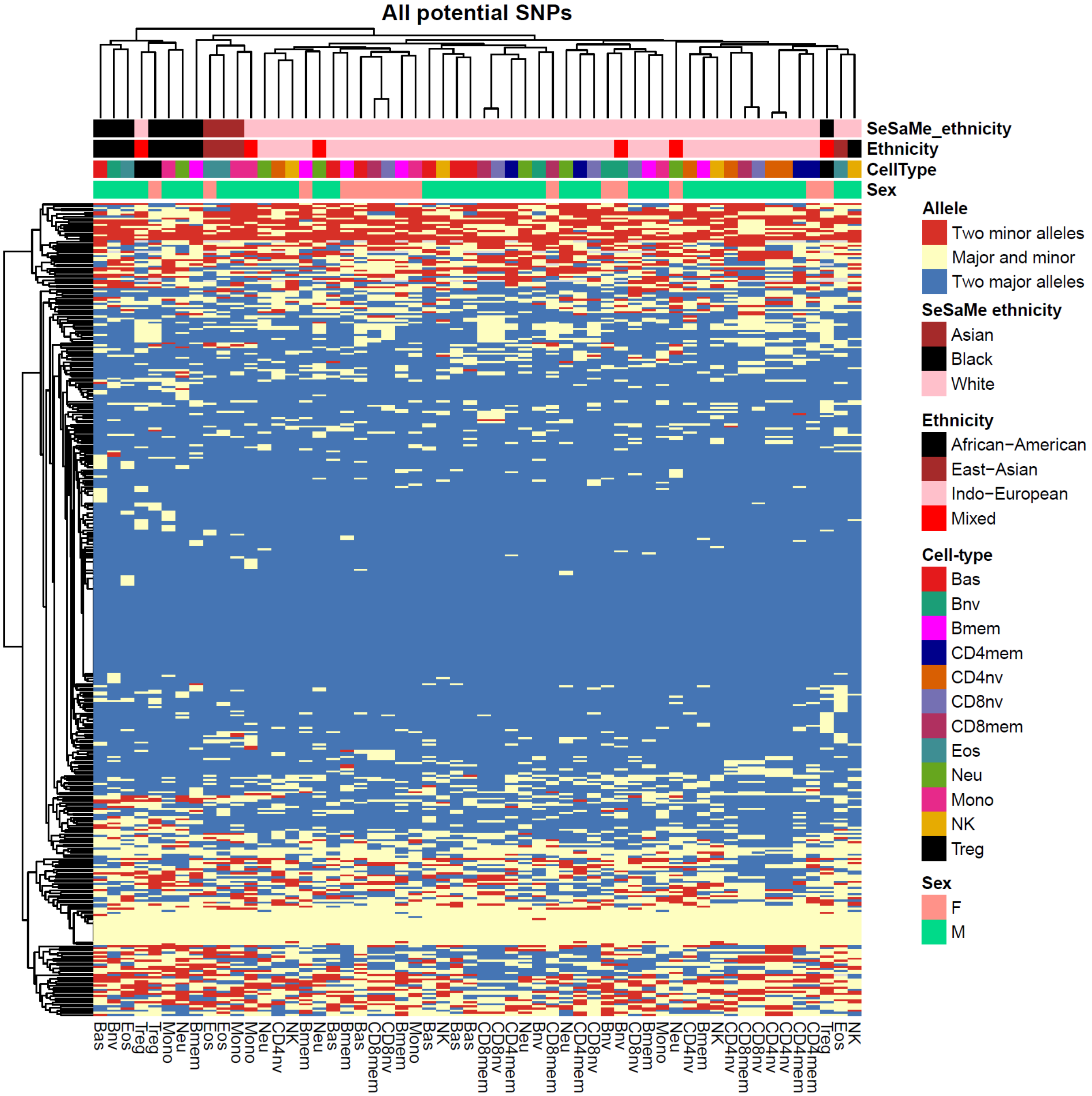

Major and minor alleles were determined using the in-band allele according to Zhou et al. Although named “ethnicity” SeSaMe uses SNP ancestry markers to group the subjects using a random forest approach, the resulting names are based on race. Ethnicity is grouped into major ancestry groups based on self-report of the donors. Here the category of African-American describes self-reported black individuals born in the USA and potentially with different unknown degrees of ancestry admixture. The Mixed category includes subjects reporting multiple ethnicities and those reported as Hispanic/Latinos and potentially with an unknown ancestry admixture.

Supplementary Fig. 3. Principal component regression analysis of phenotype and technical variables for the samples included in the library

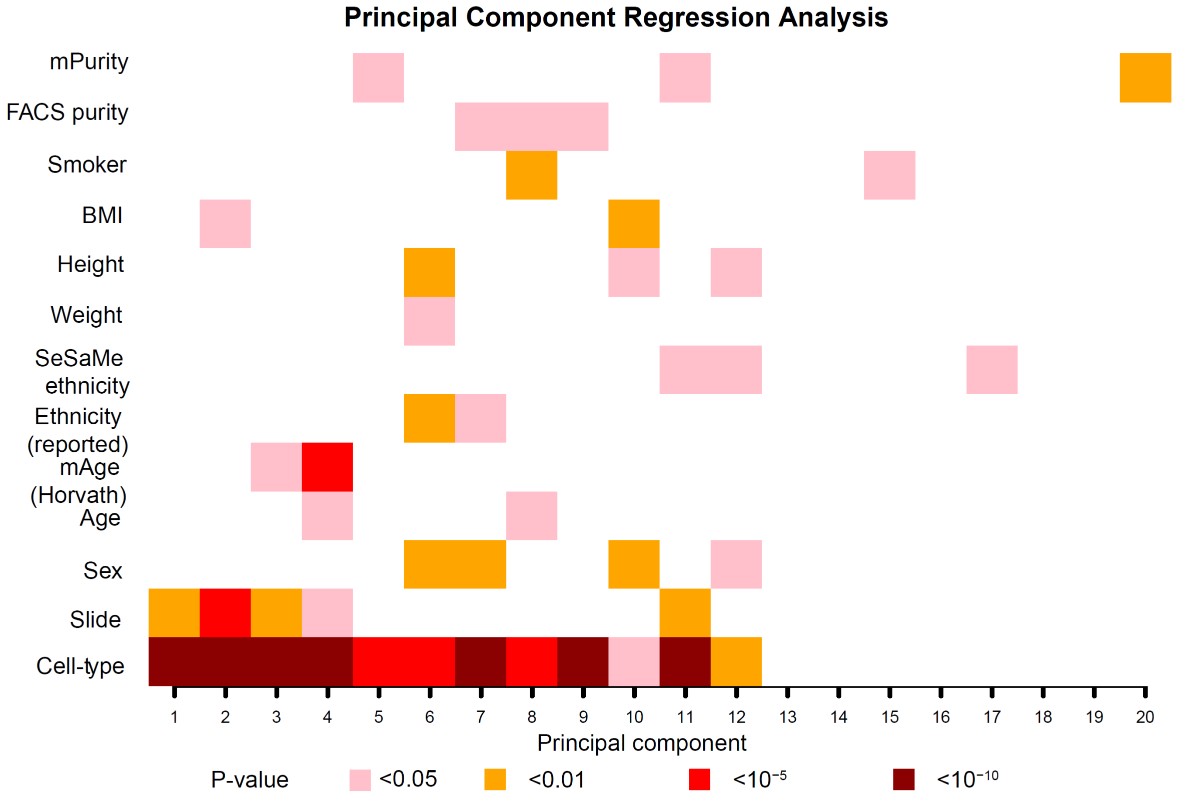

Notes: mPurity: methylation purity derived from the automatic deconvolution by Jaffe et al. FACS purity: reported flow sorting estimates of purity. BMI: body mass index (continuous). SeSaMe "ethnicity": refers to single nucleotide polymorphisms ancestry markers, and the random forest approach by Zhou et al. Ethnicity (reported) is based on self-reported ethnicity of the donor. mAge (Horvath): methylation age using Horvath's epigenetic clock.

Supplementary Table 2. Dirichlet distributions of artificial mixtures used for training and testing for IDOL.

| | Bas | Bmem | Bnv | CD4mem | CD4nv | CD8mem | CD8nv | Eos | Mono | Neu | NK | Treg |
|---|---|---|---|---|---|---|---|---|---|---|---|---|
| MIX_1 | 0 | 8.4 | 14.1 | 13 | 5.7 | 7.7 | 13.2 | 3.9 | 8.2 | 8.8 | 17 | 0 |
| MIX_2 | 0 | 13.3 | 7.7 | 7.8 | 16.8 | 5.4 | 5.9 | 20 | 6.5 | 5.8 | 10.8 | 0 |
| MIX_3 | 9.4 | 4.4 | 6.4 | 20.8 | 10.8 | 8.2 | 8 | 6.5 | 9.5 | 8.2 | 7.8 | 0 |
| MIX_4 | 10.4 | 8.7 | 23.8 | 8.9 | 8.8 | 5.5 | 5.5 | 6.5 | 5.5 | 7.5 | 8.9 | 0 |
| MIX_5 | 0 | 11.2 | 14.4 | 0 | 8 | 8.2 | 5.6 | 10.8 | 18.1 | 12 | 5.7 | 6 |
| MIX_6 | 0 | 7.5 | 11 | 0 | 9.9 | 11.6 | 8.7 | 10.6 | 5.2 | 15.4 | 9 | 11.1 |
| MIX_7 | 0 | 5.1 | 14.7 | 0 | 3.3 | 8.4 | 5.6 | 22.7 | 10.2 | 7.2 | 6.8 | 16 |
| MIX_8 | 0 | 16.3 | 9.7 | 0 | 18.4 | 6.5 | 9.3 | 12.9 | 2.2 | 7.5 | 4.7 | 12.5 |
| MIX_9 | 0 | 7.3 | 8.7 | 5.4 | 14 | 7.7 | 7.3 | 16.2 | 3.2 | 4.6 | 10.4 | 15.2 |
| MIX_10 | 0 | 16.5 | 11.3 | 8.2 | 10.5 | 9.2 | 6.2 | 5.8 | 6 | 3.7 | 12.5 | 10.1 |
| MIX_11 | 8 | 7.4 | 4.3 | 16.3 | 8.9 | 9.4 | 8.9 | 9.7 | 9.7 | 7.8 | 6 | 3.6 |
| MIX_12 | 13.2 | 3.6 | 12.4 | 11.4 | 14.8 | 3.9 | 7.3 | 8.6 | 5.3 | 6.7 | 2.8 | 10 |

Supplementary Table 3. Comparison of different sizes of optimal libraries using IDOL for EPIC technology, and the legacy IDOL in the 450k common probes

| | EPIC IDOL-Ext | | 450k IDOL-Ext | |
|---|---|---|---|---|
| | RMSE | $R^2$ | RMSE | $R^0$ |
| CpG_250 | 0.631 | 1 | 0.612 | 1 |
| CpG_300 | 0.526 | 1 | 0.505 | 1 |
| CpG_350 | 0.499 | 1 | 0.608 | 1 |
| CpG_400 | 0.404 | 1 | 0.373 | 1 |
| CpG_450 | 0.342 | 1 | 0.394 | 1 |
| CpG_500 | 0.338 | 1 | 0.255 | 1 |
| CpG_550 | 0.342 | 1 | 0.344 | 1 |
| CpG_600 | 0.307 | 1 | 0.235 | 1 |
| CpG_650 | 0.293 | 1 | 0.249 | 1 |
| CpG_700 | 0.446 | 1 | 0.233 | 1 |
| CpG_750 | 0.281 | 1 | 0.232 | 1 |
| CpG_800 | 0.323 | 1 | 0.299 | 1 |
| CpG_850 | 0.274 | 1 | 0.296 | 1 |
| CpG_900 | 0.277 | 1 | 0.295 | 1 |
| CpG_950 | 0.250 | 1 | 0.212 | 1 |
| CpG_1000 | 0.259 | 1 | 0.202 | 1 |
| CpG_1050 | 0.238 | 1 | 0.205 | 1 |
| CpG_1100 | 0.254 | 1 | 0.198 | 1 |
| CpG_1200 | **0.226** | **1** | 0.219 | 1 |
| CpG_1300 | 0.255 | 1 | 0.498 | 1 |
| CpG_1400 | 0.260 | 1 | 0.186 | 1 |
| CpG_1500 | 0.231 | 1 | **0.175** | **1** |
| CpG_1600 | 0.252 | 1 | 0.205 | 1 |
| CpG_1700 | 0.242 | 1 | 0.304 | 1 |
| CpG_1800 | 0.235 | 1 | 0.211 | 1 |
| CpG_1900 | 0.257 | 1 | 0.389 | 1 |
| CpG_2000 | 0.45 | 1 | 0.339 | 1 |
| CpG_2500 | 0.466 | 1 | 0.368 | 1 |
| CpG_3000 | 1.088 | 0.999 | 0.705 | 1 |

Notes: RMSE (root mean square error), $R^2$ (coefficient of determination). The bolded cells are the selected optimal IDOL libraries.

Supplementary Fig. 4. Comparison of the EPIC IDOL-ext, 450k IDOL-Ext and minfi pickCompProbes automatic selection estimations per cell-type. Automatic selection is severely biased for T-cells subtypes, Bcell naïve and eosinophils

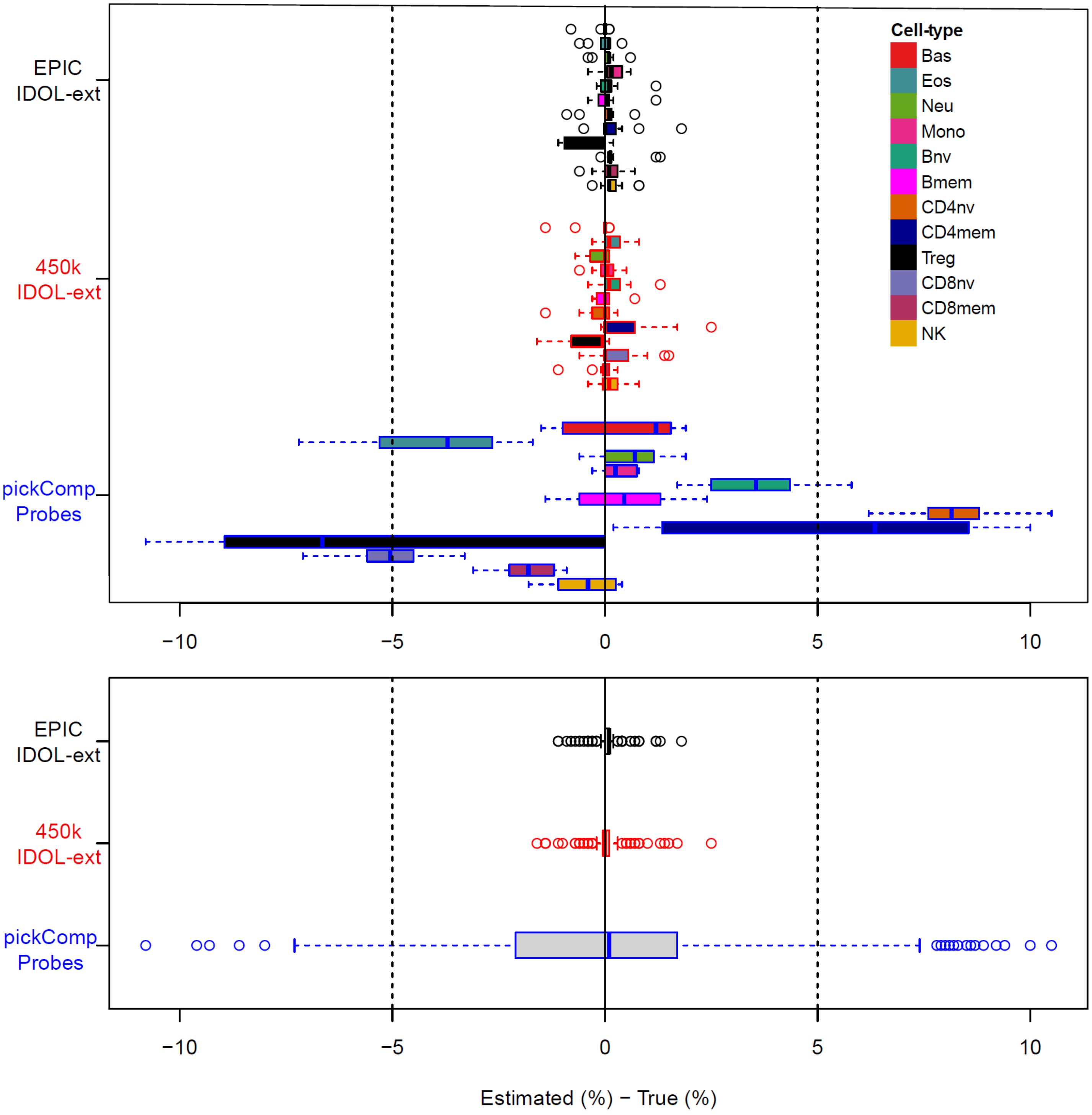

Notes: the color of the box correspond to the library selection method , for the top panel the colors inside the box correspond to the cell-type in the same order as the legend of the right side

Supplementary Tables 4, 5, 6
(see Excel file).

Supplementary Fig. 5. Comparison between cell proportion estimations, and cell counts estimations in whole blood DNA samples with FCM data.

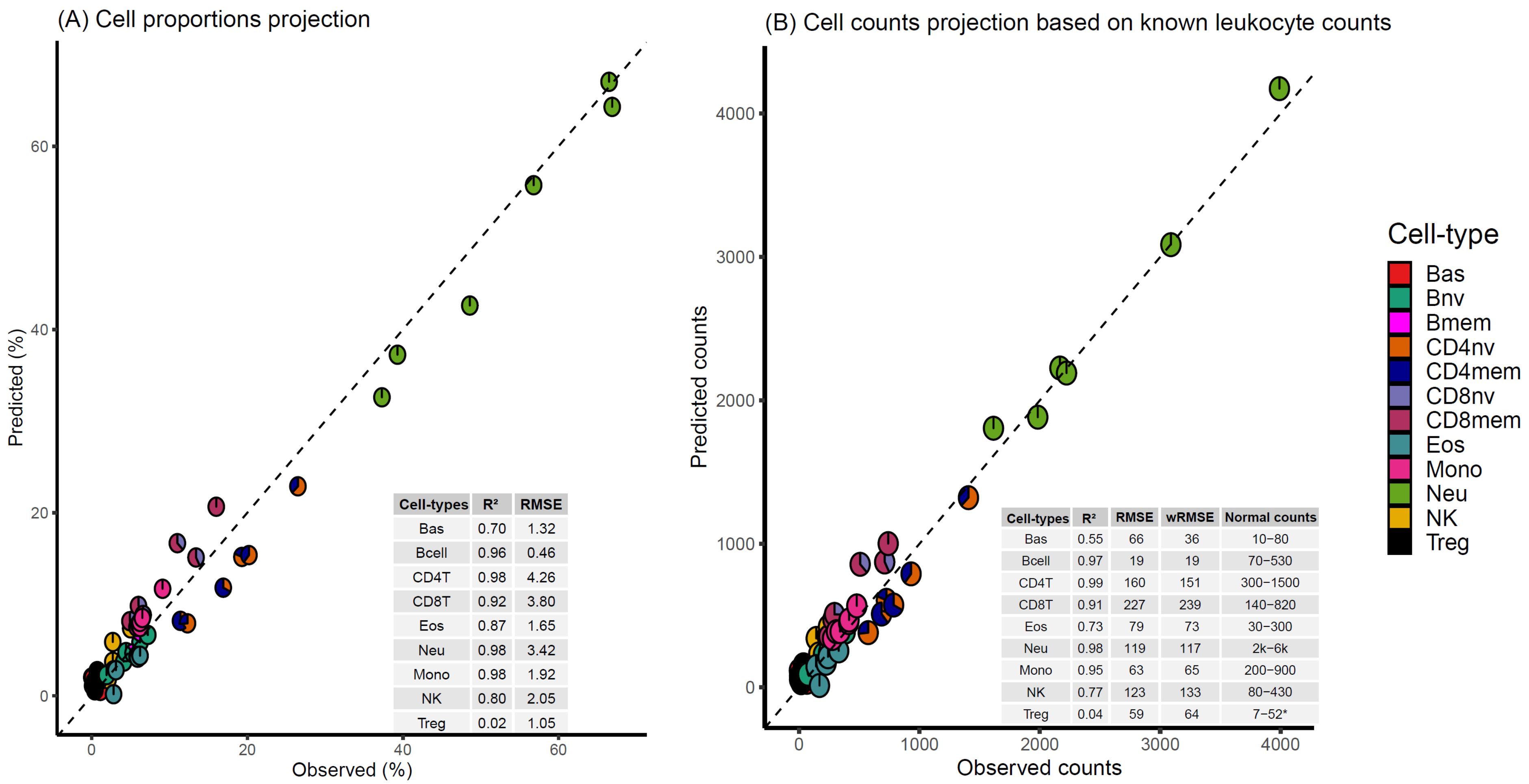

| Cell-types | R² | RMSE |
|---|---|---|
| Bas | 0.70 | 1.32 |
| Bcell | 0.96 | 0.46 |
| CD4T | 0.98 | 4.26 |
| CD8T | 0.92 | 3.80 |
| Eos | 0.87 | 1.65 |
| Neu | 0.98 | 3.42 |
| Mono | 0.98 | 1.92 |
| NK | 0.80 | 2.05 |
| Treg | 0.02 | 1.05 |

| Cell-types | R² | RMSE | wRMSE | Normal counts |
|---|---|---|---|---|
| Bas | 0.55 | 66 | 36 | 10−80 |
| Bcell | 0.97 | 19 | 19 | 70−530 |
| CD4T | 0.99 | 160 | 151 | 300−1500 |
| CD8T | 0.91 | 227 | 239 | 140−820 |
| Eos | 0.73 | 79 | 73 | 30−300 |
| Neu | 0.98 | 119 | 117 | 2k−6k |
| Mono | 0.95 | 63 | 65 | 200−900 |
| NK | 0.77 | 123 | 133 | 80−430 |
| Treg | 0.04 | 59 | 64 | 7−52* |

Notes: The area of each pieplot corresponds to the estimated proportion of the cell-types within each group. CD4Tcorresponds to the sum of CD4+ T cells naïve, memory, Treg were measured independently. CD8Tcorresponds to the sum of CD8+ T cells naïve and memory. Bcell to the sum of the naïve and memory. Panel B: a measure of weighted root mean square error was added according to the true (FCM) proportions of the measured cells. Normal counts are based on information frme STEM-cells. *The normal range of Tregs is not established so the range presented could not reflect the actual counts in peripheral blood.

Supplementary Fig. 6. Validation of additional components of the EPIC IDOL-ext (Panel A) and 450k IDOL-ext (Panel B) libraries using flow cytometry

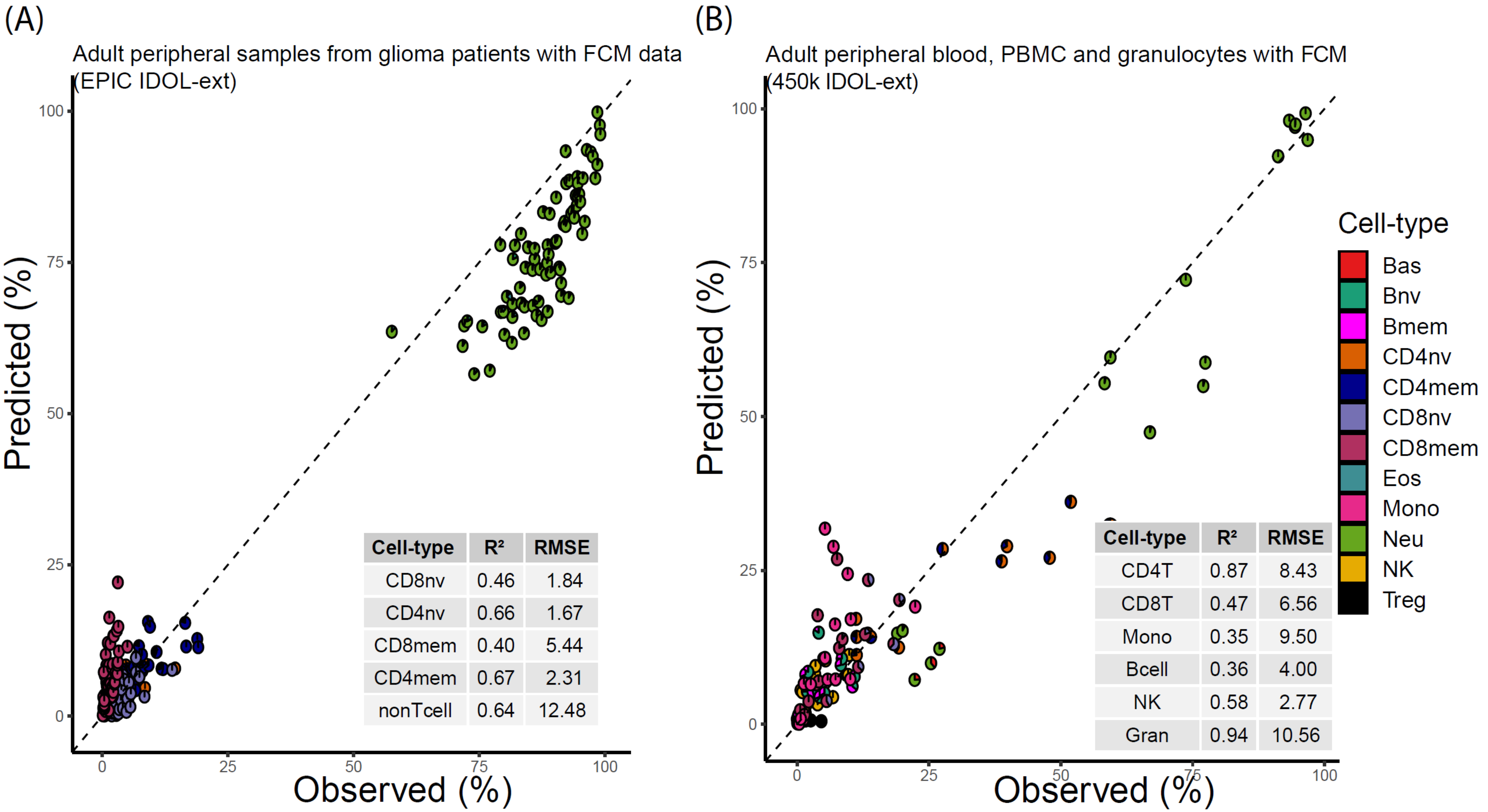

Notes: The area of each pieplot corresponds to the estimated proportion of the cell-types within each group. Panel A (EPIC data) includes information from the T-cell memory and naïve subtypes only, the unmeasured cells are summed as non-Tcells. Panel B (450k) Gran (Granulocytes) corresponds to the sum of Neu-neutrophils, Eos-eosinophils, and Bas-basophils. CD4Tcorresponds to the sum of CD4+ T cells naïve, memory and Treg. CD8T corresponds to the sum of CD8+ T cells naïve and memory. Bcell to the sum of the naïve and memory. Information in panel B was retrieved from the supplementary materials from Reinius et al. 2012 for whole blood samples, peripheral blood mononuclear cells, and granulocytes.

Supplementary Fig. 7. Exploratory analysis applying the libraries to umbilical cord blood datasets. Panel A (450K IDOL-ext): contains FCM information from umbilical cord blood samples. Panel B (EPIC IDOL-ext) corresponds to artificial mixtures using cells isolated from umbilical cord blood.

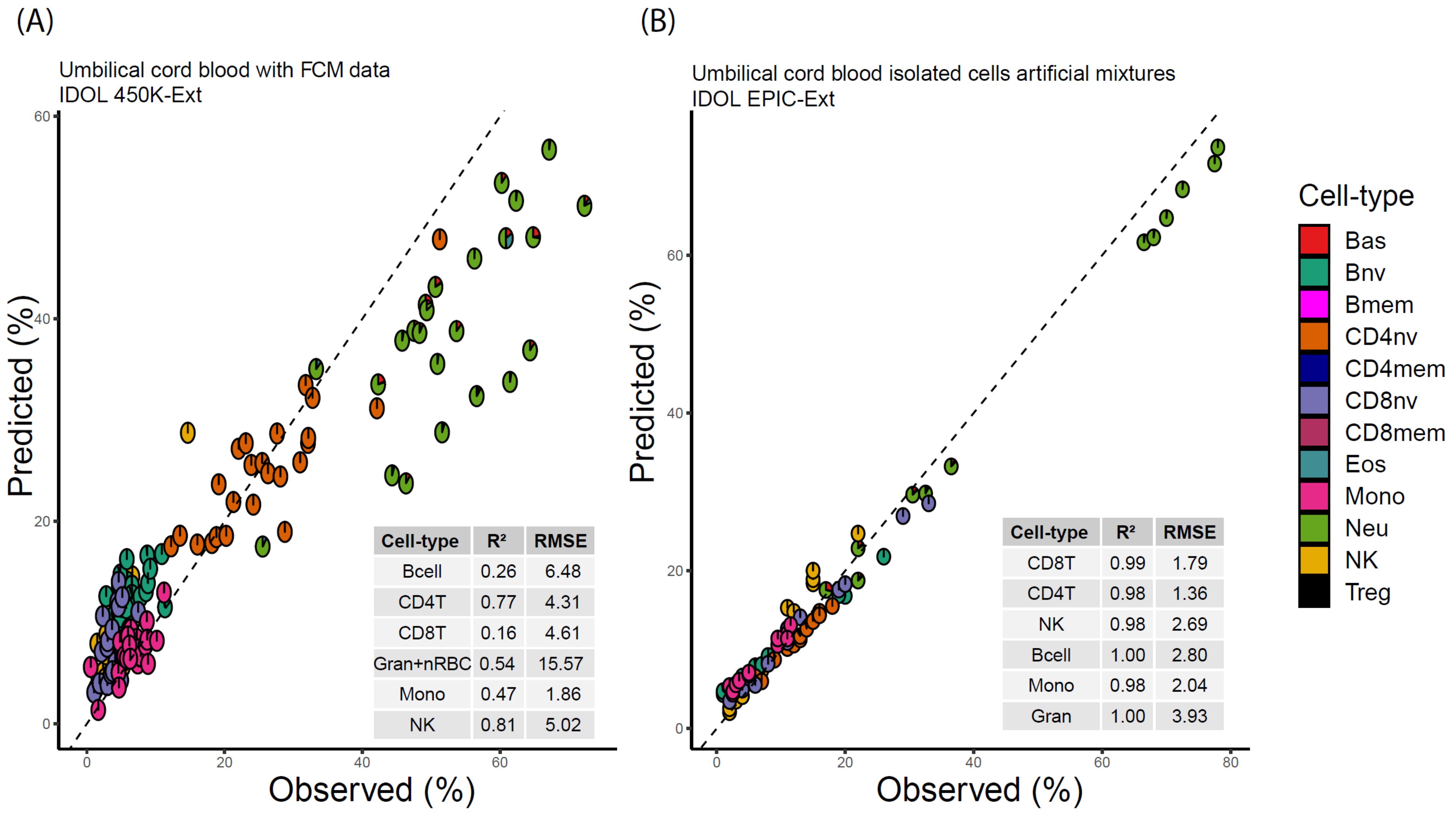

| Cell-type | $R^2$ | RMSE |
|---|---|---|
| Bcell | 0.26 | 6.48 |
| CD4T | 0.77 | 4.31 |
| CD8T | 0.16 | 4.61 |
| Gran+nRBC | 0.54 | 15.57 |
| Mono | 0.47 | 1.86 |
| NK | 0.81 | 5.02 |

| Cell-type | $R^2$ | RMSE |
|---|---|---|
| CD8T | 0.99 | 1.79 |
| CD4T | 0.98 | 1.36 |
| NK | 0.98 | 2.69 |
| Bcell | 1.00 | 2.80 |
| Mono | 0.98 | 2.04 |
| Gran | 1.00 | 3.93 |

Notes: The area of each pieplot corresponds to the estimated proportion of the cell-types within each group. Panel A (450k IDOL-ext) Gran (Granulocytes) corresponds to the sum of Neu-neutrophils, Eos-eosinophils, and Bas-basophils. CD4Tcorresponds to the sum of CD4+ T cells naïve, memory and Treg. CD8T corresponds to the sum of CD8+ T cells naïve and memory. Bcell to the sum of the naïve and memory.

Supplementary Table 7. Gene Enrichment Analyses using missMethyl and the mSigDB (GSEA) curated database v7.2

| GSEA pathway (v.7.2) | N | DM | P-value | FDR |
|---|---|---|---|---|
| AIZARANI LIVER C1 NK NKT CELLS 1 | 147 | 32 | 3.61E-12 | 1.12E-07 |
| ZHENG BOUND BY FOXP3 | 479 | 68 | 2.62E-11 | 3.62E-07 |
| GSE10325 LUPUS CD4 TCELL VS LUPUS BCELL UP | 188 | 37 | 4.66E-11 | 3.62E-07 |
| GO T CELL ACTIVATION | 455 | 55 | 4.46E-11 | 3.62E-07 |
| GO CELL ACTIVATION | 1338 | 112 | 7.33E-11 | 4.56E-07 |
| GSE10325 CD4 TCELL VS MYELOID UP | 189 | 35 | 1.65E-10 | 8.54E-07 |
| GO POSITIVE REGULATION OF IMMUNE SYSTEM PROCESS | 940 | 82 | 7.62E-10 | 3.38E-06 |
| GO REGULATION OF CELL CELL ADHESION | 419 | 52 | 1.37E-09 | 4.74E-06 |
| HAY BONE MARROW NAIVE T CELL | 372 | 46 | 1.35E-09 | 4.74E-06 |
| GO LYMPHOCYTE ACTIVATION | 652 | 64 | 2.95E-09 | 8.30E-06 |
| GO POSITIVE REGULATION OF IMMUNE RESPONSE | 621 | 60 | 3.20E-09 | 8.30E-06 |
| GO REGULATION OF T CELL ACTIVATION | 317 | 40 | 3.19E-09 | 8.30E-06 |
| GO LEUKOCYTE CELL CELL ADHESION | 343 | 42 | 5.06E-09 | 1.21E-05 |
| GSE1460 INTRATHYMIC T PROGENITOR VS CD4 THYMOCYTE DN | 191 | 34 | 5.51E-09 | 1.22E-05 |
| CHEN METABOLIC SYNDROM NETWORK | 1180 | 101 | 9.40E-09 | 1.86E-05 |
| GO REGULATION OF IMMUNE SYSTEM PROCESS | 1443 | 108 | 9.59E-09 | 1.86E-05 |
| GO REGULATION OF CELL ADHESION | 701 | 74 | 1.07E-08 | 1.96E-05 |
| GO REGULATION OF IMMUNE RESPONSE | 857 | 71 | 1.45E-08 | 2.37E-05 |
| GO REGULATION OF CELL ACTIVATION | 553 | 55 | 1.38E-08 | 2.37E-05 |
| LEE DIFFERENTIATING T LYMPHOCYTE | 189 | 32 | 1.73E-08 | 2.69E-05 |
| GSE4590 LARGE PRE BCELL VS VPREB POS LARGE PRE BCELL DN | 174 | 32 | 2.21E-08 | 3.27E-05 |
| GSE10325 CD4 TCELL VS BCELL UP | 184 | 32 | 2.52E-08 | 3.56E-05 |
| GO REGULATION OF LYMPHOCYTE ACTIVATION | 422 | 45 | 1.23E-07 | 1.67E-04 |
| GO NEGATIVE REGULATION OF CELL CELL ADHESION | 181 | 26 | 3.05E-07 | 3.95E-04 |
| GSE11057 NAIVE VS CENT MEMORY CD4 TCELL DN | 193 | 32 | 3.49E-07 | 4.27E-04 |
| GSE22886 NAIVE TCELL VS MONOCYTE UP | 187 | 29 | 3.98E-07 | 4.27E-04 |
| GSE22886 NAIVE CD4 TCELL VS MONOCYTE UP | 181 | 29 | 3.67E-07 | 4.27E-04 |
| MANNO MIDBRAIN NEUROTYPES HMGL | 552 | 53 | 3.86E-07 | 4.27E-04 |
| HAY BONE MARROW NK CELLS | 346 | 41 | 3.93E-07 | 4.27E-04 |
| HU FETAL RETINA MICROGLIA | 357 | 40 | 4.59E-07 | 4.76E-04 |
| GO POSITIVE REGULATION OF LEUKOCYTE CELL CELL ADHESION | 222 | 30 | 5.30E-07 | 5.32E-04 |
| GO CD4 POSITIVE ALPHA BETA T CELL ACTIVATION | 91 | 17 | 6.17E-07 | 5.99E-04 |
| MARTENS BOUND BY PML RARA FUSION | 447 | 53 | 9.18E-07 | 8.65E-04 |
| GSE11386 NAIVE VS MEMORY BCELL DN | 151 | 27 | 1.25E-06 | 1.14E-03 |
| GSE14699 NAIVE VS DELETIONAL TOLERANCE CD8 TCELL UP | 147 | 26 | 1.30E-06 | 1.16E-03 |
| GSE22886 TCELL VS BCELL NAIVE UP | 193 | 27 | 1.45E-06 | 1.16E-03 |
| GSE22886 NAIVE CD8 TCELL VS DC UP | 186 | 26 | 1.37E-06 | 1.16E-03 |
| GSE15330 LYMPHOID MULTIPOTENT VS MEGAKARYOCYTE ERYTHROID PROGENITOR IKAROS KO DN | 169 | 25 | 1.47E-06 | 1.16E-03 |
| GO ACTIVATION OF IMMUNE RESPONSE | 428 | 44 | 1.46E-06 | 1.16E-03 |
| GO CELL ACTIVATION INVOLVED IN IMMUNE RESPONSE | 683 | 56 | 1.49E-06 | 1.16E-03 |
| GSE7460 CD8 TCELL VS CD4 TCELL ACT UP | 185 | 26 | 1.53E-06 | 1.16E-03 |
| GO CD4 POSITIVE ALPHA BETA T CELL DIFFERENTIATION | 75 | 15 | 1.70E-06 | 1.24E-03 |
| GO POSITIVE REGULATION OF CELL ACTIVATION | 330 | 36 | 1.72E-06 | 1.24E-03 |

| GSEA pathway (v.7.2) | N | DM | P-value | FDR |
|---|---|---|---|---|
| RYTTCCTG ETS2 B | 1052 | 90 | 1.96E-06 | 1.39E-03 |
| AIZARANI LIVER C5 NK NKT CELLS 3 | 115 | 19 | 2.26E-06 | 1.56E-03 |
| GSE13306 LAMINA PROPRIA VS SPLEEN TREG UP | 192 | 27 | 2.36E-06 | 1.58E-03 |
| GO IMMUNE EFFECTOR PROCESS | 1131 | 80 | 2.53E-06 | 1.58E-03 |
| GO ALPHA BETA T CELL ACTIVATION | 137 | 21 | 2.46E-06 | 1.58E-03 |
| GO POSITIVE REGULATION OF LYMPHOCYTE ACTIVATION | 269 | 31 | 2.54E-06 | 1.58E-03 |
| FAN EMBRYONIC CTX BRAIN NAIVE LIKE T CELL | 152 | 22 | 2.41E-06 | 1.58E-03 |
| GO T CELL DIFFERENTIATION | 247 | 30 | 2.70E-06 | 1.61E-03 |
| GAO LARGE INTESTINE ADULT CI MESENCHYMAL CELLS | 338 | 39 | 2.70E-06 | 1.61E-03 |
| GSE10325 LUPUS CD4 TCELL VS LUPUS MYELOID UP | 182 | 27 | 3.19E-06 | 1.82E-03 |
| GSE7218 UNSTIM VS ANTIGEN STIM THROUGH IGG BCELL DN | 165 | 22 | 3.11E-06 | 1.82E-03 |
| GSE24726 WT VS E2 2 KO PDC DAY6 POST DELETION DN | 187 | 28 | 3.21E-06 | 1.82E-03 |
| GSE29618 BCELL VS PDC UP | 178 | 26 | 3.37E-06 | 1.85E-03 |
| GO NEGATIVE REGULATION OF LEUKOCYTE CELL CELL ADHESION | 128 | 18 | 3.38E-06 | 1.85E-03 |
| GO ALPHA BETA T CELL DIFFERENTIATION | 102 | 18 | 4.23E-06 | 2.27E-03 |
| GSE22886 DC VS MONOCYTE DN | 190 | 26 | 4.53E-06 | 2.27E-03 |
| GSE37301 PRO BCELL VS CD4 TCELL UP | 161 | 25 | 4.50E-06 | 2.27E-03 |
| GAO LARGE INTESTINE 24W C11 PANETH LIKE CELL | 307 | 33 | 4.39E-06 | 2.27E-03 |
| CUI DEVELOPING HEART C9 B T CELL | 147 | 22 | 4.41E-06 | 2.27E-03 |
| GO IMMUNE RESPONSE REGULATING SIGNALING PATHWAY | 382 | 41 | 6.34E-06 | 3.13E-03 |
| GO POSITIVE REGULATION OF CELL CELL ADHESION | 266 | 32 | 6.71E-06 | 3.26E-03 |
| GO REGULATORY T CELL DIFFERENTIATION | 34 | 9 | 6.85E-06 | 3.28E-03 |
| GO CD8 POSITIVE ALPHA BETA T CELL DIFFERENTIATION | 15 | 7 | 6.96E-06 | 3.28E-03 |
| GSE17186 NAIVE VS CD21LOW TRANSITIONAL BCELL CORD BLOOD UP | 189 | 27 | 8.17E-06 | 3.63E-03 |
| GSE16266 CTRL VS LPS STIM MEF UP | 190 | 28 | 8.13E-06 | 3.63E-03 |
| GO POSITIVE REGULATION OF CELL ADHESION | 410 | 45 | 8.05E-06 | 3.63E-03 |
| ZHENG CORD BLOOD C10 MULTILYMPHOID PROGENITOR | 91 | 18 | 7.85E-06 | 3.63E-03 |
| GSE25087 TREG VS TCONV ADULT UP | 194 | 26 | 8.55E-06 | 3.75E-03 |
| GSE37301 HEMATOPOIETIC STEM CELL VS GRAN MONO PROGENITOR UP | 156 | 24 | 8.76E-06 | 3.75E-03 |
| GO LEUKOCYTE PROLIFERATION | 290 | 29 | 8.81E-06 | 3.75E-03 |
| GO NEGATIVE REGULATION OF IMMUNE SYSTEM PROCESS | 439 | 40 | 9.67E-06 | 4.06E-03 |
| GSE22601 DOUBLE POSITIVE VS CD8 SINGLE POSITIVE THYMOCYTE UP | 198 | 27 | 1.00E-05 | 4.13E-03 |
| GSE3039 ALPHABETA CD8 TCELL VS B1 BCELL DN | 187 | 24 | 1.01E-05 | 4.13E-03 |
| GSE14415 NATURAL TREG VS TCONV UP | 153 | 22 | 1.08E-05 | 4.30E-03 |
| GO NEGATIVE REGULATION OF LYMPHOCYTE ACTIVATION | 145 | 20 | 1.08E-05 | 4.30E-03 |
| GSE36476 CTRL VS TSST ACT 16H MEMORY CD4 TCELL YOUNG UP | 188 | 26 | 1.25E-05 | 4.85E-03 |
| REACTOME INNATE IMMUNE SYSTEM | 986 | 71 | 1.24E-05 | 4.85E-03 |
| GSE29949 MICROGLIA BRAIN VS CD8 POS DC SPLEEN UP | 186 | 25 | 1.31E-05 | 5.02E-03 |
| GO REGULATION OF ANTIGEN RECEPTOR MEDIATED SIGNALING PATHWAY | 66 | 13 | 1.35E-05 | 5.13E-03 |
| GSE14415 INDUCED TREG VS FOXP3 KO INDUCED TREG UP | 161 | 23 | 1.40E-05 | 5.23E-03 |
| GSE22886 NAIVE TCELL VS DC UP | 191 | 24 | 1.43E-05 | 5.27E-03 |
| GSE13522 WT VS IFNAR KO SKING T CRUZI Y STRAIN INF UP | 196 | 27 | 1.44E-05 | 5.27E-03 |
| GSE16385 ROSIGLITAZONE IL4 VS IFNG TNF STIM MACROPHAGE UP | 192 | 27 | 1.47E-05 | 5.30E-03 |
| ZNF597 TARGET GENES | 769 | 63 | 1.48E-05 | 5.30E-03 |
| GSE22886 NAIVE CD8 TCELL VS MONOCYTE UP | 181 | 25 | 1.52E-05 | 5.36E-03 |
| GSE36476 CTRL VS TSST ACT 40H MEMORY CD4 TCELL YOUNG UP | 193 | 26 | 1.70E-05 | 5.95E-03 |

| GSEA pathway (v.7.2) | N | DM | P-value | FDR |
|---|---|---|---|---|
| DURANTE ADULT OLFACTORY NEUROEPITHELIUM NK CELLS | 78 | 13 | 1.86E-05 | 6.41E-03 |
| DURANTE ADULT OLFACTORY NEUROEPITHELIUM CD8 T CELLS | 64 | 11 | 1.89E-05 | 6.45E-03 |
| ZHENG FOXP3 TARGETS IN T LYMPHOCYTE DN | 39 | 12 | 1.91E-05 | 6.46E-03 |
| GSE11057 NAIVE VS EFF MEMORY CD4 TCELL UP | 185 | 27 | 2.00E-05 | 6.68E-03 |
| HAY BONE MARROW CD8 T CELL | 67 | 12 | 2.07E-05 | 6.86E-03 |
| GSE29618 BCELL VS MONOCYTE UP | 175 | 24 | 2.23E-05 | 7.30E-03 |
| GSE14699 NAIVE VS ACT CD8 TCELL DN | 167 | 25 | 2.33E-05 | 7.54E-03 |
| GO NEGATIVE REGULATION OF CELL ADHESION | 285 | 32 | 2.38E-05 | 7.63E-03 |
| GSE13547 CTRL VS ANTI IGM STIM ZFX KO BCELL 12H DN | 153 | 24 | 2.62E-05 | 8.32E-03 |
| GSE14350 TREG VS TEFF UP | 191 | 25 | 2.68E-05 | 8.34E-03 |
| GSE9988 ANTI TREM1 VS VEHICLE TREATED MONOCYTES DN | 193 | 26 | 2.68E-05 | 8.34E-03 |
| GSE22886 NAIVE BCELL VS MONOCYTE DN | 196 | 26 | 2.73E-05 | 8.41E-03 |
| GSE26343 WT VS NFAT5 KO MACROPHAGE LPS STIM UP | 197 | 23 | 2.90E-05 | 8.52E-03 |
| GO CD8 POSITIVE ALPHA BETA T CELL ACTIVATION | 26 | 8 | 2.84E-05 | 8.52E-03 |
| GO B CELL RECEPTOR SIGNALING PATHWAY | 60 | 13 | 2.87E-05 | 8.52E-03 |
| ZNF184 TARGET GENES | 1283 | 93 | 2.90E-05 | 8.52E-03 |
| CUI DEVELOPING HEART C8 MACROPHAGE | 264 | 28 | 2.84E-05 | 8.52E-03 |
| HADDAD B LYMPHOCYTE PROGENITOR | 272 | 33 | 3.46E-05 | 1.00E-02 |
| GSE3565 CTRL VS LPS INJECTED SPLENOCYTES DN | 169 | 24 | 3.47E-05 | 1.00E-02 |
| SIG INSULIN RECEPTOR PATHWAY IN CARDIAC MYOCYTES | 48 | 12 | 3.63E-05 | 1.04E-02 |
| AIZARANI LIVER C3 NK NKT CELLS 2 | 149 | 20 | 3.68E-05 | 1.04E-02 |
| THEILGAARD NEUTROPHIL AT SKIN WOUND DN | 216 | 28 | 3.89E-05 | 1.08E-02 |
| GO LYMPHOCYTE DIFFERENTIATION | 347 | 34 | 3.88E-05 | 1.08E-02 |
| GSE13306 RA VS UNTREATED MEM CD4 TCELL UP | 194 | 26 | 3.94E-05 | 1.09E-02 |
| GSE7219 UNSTIM VS LPS AND ANTI CD40 STIM NIK NFKB2 KO DC DN | 190 | 25 | 4.17E-05 | 1.14E-02 |
| GOLDRATH NAIVE VS EFF CD8 TCELL UP | 194 | 25 | 4.56E-05 | 1.23E-02 |
| GSE31082 DN VS CD8 SP THYMOCYTE DN | 196 | 27 | 4.66E-05 | 1.23E-02 |
| GSE26488 CTRL VS PEPTIDE INJECTION OT2 THYMOCYTE DN | 178 | 25 | 4.61E-05 | 1.23E-02 |
| GO CELL CELL ADHESION | 840 | 71 | 4.66E-05 | 1.23E-02 |
| GNF2 JAK1 | 32 | 9 | 4.78E-05 | 1.25E-02 |
| GSE16450 IMMATURE VS MATURE NEURON CELL LINE UP | 181 | 25 | 4.86E-05 | 1.26E-02 |
| GSE36476 CTRL VS TSST ACT 72H MEMORY CD4 TCELL YOUNG UP | 190 | 25 | 5.04E-05 | 1.30E-02 |
| GSE17721 POLYIC VS PAM3CSK4 16H BMDC UP | 195 | 24 | 5.14E-05 | 1.30E-02 |
| GSE17974 CTRL VS ACT IL4 AND ANTI IL12 24H CD4 TCELL UP | 182 | 24 | 5.25E-05 | 1.30E-02 |
| GSE23502 BM VS COLON TUMOR MYELOID DERIVED SUPPRESSOR CELL DN | 192 | 23 | 5.23E-05 | 1.30E-02 |
| GSE33425 CD8 ALPHAALPHA VS ALPHABETA CD161 HIGH TCELL DN | 196 | 23 | 5.17E-05 | 1.30E-02 |
| ZHONG PFC MAJOR TYPES MICROGLIA | 414 | 40 | 5.26E-05 | 1.30E-02 |
| GSE9988 ANTI TREM1 VS CTRL TREATED MONOCYTES DN | 197 | 25 | 5.91E-05 | 1.45E-02 |
| OSMAN BLADDER CANCER DN | 394 | 37 | 6.07E-05 | 1.45E-02 |
| MULLIGHAN MLL SIGNATURE 1 UP | 365 | 35 | 6.02E-05 | 1.45E-02 |
| GSE17721 LPS VS CPG 1H BMDC DN | 197 | 24 | 6.05E-05 | 1.45E-02 |
| GO REGULATION OF LEUKOCYTE PROLIFERATION | 227 | 23 | 6.18E-05 | 1.47E-02 |
| GO 14 3 3 PROTEIN BINDING | 31 | 10 | 6.41E-05 | 1.51E-02 |
| GSE15330 HSC VS LYMPHOID PRIMED MULTIPOTENT PROGENITOR DN | 175 | 21 | 6.49E-05 | 1.52E-02 |
| GO NEGATIVE REGULATION OF CELL ACTIVATION | 197 | 23 | 6.64E-05 | 1.54E-02 |

| GSEA pathway (v.7.2) | N | DM | P-value | FDR |
|---|---|---|---|---|
| GSE9601 UNTREATED VS NFKB INHIBITOR TREATED HCMV INF MONOCYTE DN | 169 | 23 | 6.75E-05 | 1.55E-02 |
| GSE15330 LYMPHOID MULTIPOTENT VS MEGAKARYOCYTE ERYTHROID PROGENITOR IKAROS KO UP | 158 | 23 | 6.77E-05 | 1.55E-02 |
| HALLMARK IL2 STAT5 SIGNALING | 194 | 24 | 6.92E-05 | 1.57E-02 |
| GSE45739 UNSTIM VS ACD3 ACD28 STIM NRAS KO CD4 TCELL UP | 183 | 24 | 7.33E-05 | 1.65E-02 |
| GSE21033 CTRL VS POLYIC STIM DC 12H UP | 152 | 20 | 7.45E-05 | 1.67E-02 |
| GSE8384 CTRL VS B ABORTUS 4H MAC CELL LINE DN | 189 | 22 | 8.13E-05 | 1.81E-02 |
| GSE41867 DAY6 EFFECTOR VS DAY30 EXHAUSTED CD8 TCELL LCMV CLONE13 UP | 195 | 23 | 8.34E-05 | 1.84E-02 |
| GSE26290 CTRL VS AKT INHIBITOR TREATED ANTI CD3 AND IL2 STIM CD8 TCELL DN | 189 | 25 | 8.45E-05 | 1.85E-02 |
| GSE15735 2H VS 12H HDAC INHIBITOR TREATED CD4 TCELL DN | 192 | 24 | 8.60E-05 | 1.87E-02 |
| GSE3920 IFNA VS IFNB TREATED ENDOTHELIAL CELL DN | 160 | 23 | 8.80E-05 | 1.88E-02 |
| GSE17301 ACD3 ACD28 VS ACD3 ACD28 AND IFNA5 STIM CD8 TCELL DN | 194 | 23 | 8.81E-05 | 1.88E-02 |
| GO LYMPHOCYTE ACTIVATION INVOLVED IN IMMUNE RESPONSE | 172 | 19 | 8.79E-05 | 1.88E-02 |
| GO T CELL PROLIFERATION | 182 | 20 | 8.89E-05 | 1.88E-02 |
| FAN EMBRYONIC CTX BRAIN EFFECTOR T CELL | 125 | 16 | 9.20E-05 | 1.93E-02 |
| GSE15330 HSC VS MEGAKARYOCYTE ERYTHROID PROGENITOR UP | 158 | 23 | 9.29E-05 | 1.94E-02 |
| GSE3565 DUSP1 VS WT SPLENOCYTES DN | 175 | 23 | 9.52E-05 | 1.97E-02 |
| GSE24634 NAIVE CD4 TCELL VS DAY3 IL4 CONV TREG UP | 189 | 23 | 9.71E-05 | 1.99E-02 |
| GSE18893 TCONV VS TREG 24H CULTURE DN | 186 | 23 | 9.84E-05 | 1.99E-02 |
| GSE21360 PRIMARY VS TERTIARY MEMORY CD8 TCELL UP | 190 | 23 | 9.78E-05 | 1.99E-02 |
| GSE43863 DAY6 EFF VS DAY150 MEM LY6C INT CXCR5POS CD4 TCELL UP | 191 | 25 | 9.73E-05 | 1.99E-02 |
| ZHENG FOXP3 TARGETS IN THYMUS UP | 195 | 26 | 1.02E-04 | 2.00E-02 |
| RUTELLA RESPONSE TO HGF DN | 227 | 27 | 1.02E-04 | 2.00E-02 |
| GSE1460 INTRATHYMIC T PROGENITOR VS NAIVE CD4 TCELL ADULT BLOOD DN | 185 | 23 | 1.01E-04 | 2.00E-02 |
| GSE33425 CD161 HIGH VS INT CD8 TCELL DN | 192 | 22 | 1.01E-04 | 2.00E-02 |
| GO ANTIGEN RECEPTOR MEDIATED SIGNALING PATHWAY | 232 | 27 | 1.01E-04 | 2.00E-02 |
| GSE28737 BCL6 HET VS BCL6 KO FOLLICULAR BCELL UP | 190 | 24 | 1.04E-04 | 2.03E-02 |
| GSE25677 MPL VS R848 STIM BCELL DN | 174 | 24 | 1.11E-04 | 2.15E-02 |
| KRIGE RESPONSE TO TOSEDOSTAT 24HR UP | 719 | 59 | 1.13E-04 | 2.17E-02 |
| MODULE 84 | 511 | 42 | 1.16E-04 | 2.21E-02 |
| FAN EMBRYONIC CTX BIG GROUPS MICROGLIA | 354 | 33 | 1.17E-04 | 2.22E-02 |
| POU2AF1 TARGET GENES | 788 | 61 | 1.18E-04 | 2.22E-02 |
| GSE5542 UNTREATED VS IFNA TREATED EPITHELIAL CELLS 24H UP | 192 | 28 | 1.20E-04 | 2.24E-02 |
| GOLDRATH NAIVE VS MEMORY CD8 TCELL UP | 195 | 25 | 1.24E-04 | 2.31E-02 |
| GSE3039 NKT CELL VS ALPHAALPHA CD8 TCELL DN | 191 | 21 | 1.34E-04 | 2.47E-02 |
| GSE8621 LPS STIM VS LPS PRIMED AND LPS STIM MACROPHAGE DN | 194 | 23 | 1.34E-04 | 2.47E-02 |
| GSE24574 BCL6 HIGH TFH VS TFH CD4 TCELL DN | 191 | 23 | 1.39E-04 | 2.54E-02 |
| GSE3920 UNTREATED VS IFNA TREATED FIBROBLAST UP | 165 | 22 | 1.41E-04 | 2.56E-02 |
| GSE11057 NAIVE VS MEMORY CD4 TCELL UP | 181 | 23 | 1.45E-04 | 2.61E-02 |
| GSE21063 CTRL VS ANTI IGM STIM BCELL NFATC1 KO 16H UP | 193 | 22 | 1.45E-04 | 2.61E-02 |
| MAML1 TARGET GENES | 255 | 28 | 1.50E-04 | 2.68E-02 |
| GNF2 PTPRC | 65 | 12 | 1.52E-04 | 2.70E-02 |
| GSE3982 DC VS TH1 DN | 187 | 22 | 1.53E-04 | 2.71E-02 |

| GSEA pathway (v.7.2) | N | DM | P-value | FDR |
|---|---|---|---|---|
| GSE6092 B BURGDOFERI VS B BURGDORFERI AND IFNG STIM ENDOTHELIAL CELL UP | 170 | 23 | 1.56E-04 | 2.73E-02 |
| GSE16450 IMMATURE VS MATURE NEURON CELL LINE 12H IFNA STIM UP | 192 | 25 | 1.56E-04 | 2.73E-02 |
| GSE29164 CD8 TCELL VS CD8 TCELL AND IL12 TREATED MELANOMA DAY3 DN | 191 | 23 | 1.59E-04 | 2.75E-02 |
| GSE40225 WT VS RIP B7X DIABETIC MOUSE PANCREATIC CD8 TCELL UP | 190 | 24 | 1.59E-04 | 2.75E-02 |
| JAATINEN HEMATOPOIETIC STEM CELL DN | 223 | 22 | 1.61E-04 | 2.76E-02 |
| GSE31082 DP VS CD8 SP THYMOCYTE DN | 193 | 22 | 1.64E-04 | 2.79E-02 |
| GO REGULATION OF HEMOPOIESIS | 476 | 40 | 1.64E-04 | 2.79E-02 |
| GSE22601 IMMATURE CD4 SINGLE POSITIVE VS CD8 SINGLE POSITIVE THYMOCYTE UP | 194 | 22 | 1.69E-04 | 2.85E-02 |
| GSE2770 TGFB AND IL4 ACT VS ACT CD4 TCELL 48H DN | 189 | 21 | 1.70E-04 | 2.86E-02 |
| GO T CELL ACTIVATION INVOLVED IN IMMUNE RESPONSE | 99 | 13 | 1.73E-04 | 2.90E-02 |
| KAECH NAIVE VS DAY8 EFF CD8 TCELL UP | 196 | 25 | 1.77E-04 | 2.94E-02 |
| GSE16385 IFNG TNF VS UNSTIM MACROPHAGE ROSIGLITAZONE TREATED DN | 185 | 22 | 1.81E-04 | 2.99E-02 |
| GSE17974 0H VS 2H IN VITRO ACT CD4 TCELL UP | 184 | 21 | 1.86E-04 | 3.06E-02 |
| GO REGULATION OF CD8 POSITIVE ALPHA BETA T CELL DIFFERENTIATION | 6 | 4 | 1.89E-04 | 3.09E-02 |
| GSE29618 MONOCYTE VS PDC UP | 191 | 22 | 1.92E-04 | 3.11E-02 |
| GO REGULATION OF CD4 POSITIVE ALPHA BETA T CELL ACTIVATION | 58 | 10 | 1.92E-04 | 3.11E-02 |
| ETS Q4 | 238 | 26 | 1.96E-04 | 3.16E-02 |
| GSE41867 DAY8 VS DAY15 LCMV CLONE13 EFFECTOR CD8 TCELL UP | 194 | 23 | 1.98E-04 | 3.17E-02 |
| GSE12366 PLASMA CELL VS NAIVE BCELL DN | 186 | 22 | 2.02E-04 | 3.22E-02 |
| GO NEGATIVE REGULATION OF SIGNALING | 1420 | 100 | 2.04E-04 | 3.24E-02 |
| GSE24574 NAIVE VS TCONV CD4 TCELL UP | 192 | 23 | 2.07E-04 | 3.26E-02 |
| MODULE 46 | 373 | 29 | 2.10E-04 | 3.29E-02 |
| GSE5589 LPS VS LPS AND IL10 STIM MACROPHAGE 45MIN UP | 189 | 21 | 2.10E-04 | 3.29E-02 |
| GO ACTIVATION OF MAPKKK ACTIVITY | 12 | 5 | 2.13E-04 | 3.31E-02 |
| GSE26495 NAIVE VS PD1HIGH CD8 TCELL UP | 177 | 23 | 2.14E-04 | 3.31E-02 |
| HP ACUTE LEUKEMIA | 86 | 13 | 2.19E-04 | 3.37E-02 |
| GSE9988 ANTI TREM1 AND LPS VS CTRL TREATED MONOCYTES DN | 194 | 22 | 2.24E-04 | 3.43E-02 |
| GSE16451 CTRL VS WEST EQUINE ENC VIRUS MATURE NEURON CELL LINE DN | 192 | 23 | 2.25E-04 | 3.43E-02 |
| QI PLASMACYTOMA UP | 244 | 25 | 2.35E-04 | 3.54E-02 |
| STTTCRNTTT IRF Q6 | 182 | 23 | 2.34E-04 | 3.54E-02 |
| GSE6259 33D1 POS VS DEC205 POS FLT3L INDUCED SPLENIC DC DN | 167 | 22 | 2.44E-04 | 3.66E-02 |
| RYAN MANTLE CELL LYMPHOMA NOTCH DIRECT UP | 146 | 21 | 2.45E-04 | 3.66E-02 |
| GSE21670 UNTREATED VS IL6 TREATED STAT3 KO CD4 TCELL UP | 186 | 23 | 2.46E-04 | 3.66E-02 |
| GSE40273 EOS KO VS WT TREG DN | 196 | 24 | 2.48E-04 | 3.67E-02 |
| GO ACTIN CYTOSKELETON | 490 | 49 | 2.58E-04 | 3.80E-02 |
| GO IMMUNE SYSTEM DEVELOPMENT | 976 | 71 | 2.59E-04 | 3.80E-02 |
| GO REGULATION OF LEUKOCYTE MIGRATION | 199 | 20 | 2.68E-04 | 3.90E-02 |
| GO NEGATIVE REGULATION OF REGULATORY T CELL DIFFERENTIATION | 7 | 3 | 2.68E-04 | 3.90E-02 |
| GSE21033 CTRL VS POLYIC STIM DC 24H UP | 150 | 21 | 2.70E-04 | 3.91E-02 |
| GO REGULATION OF T CELL DIFFERENTIATION | 147 | 18 | 2.72E-04 | 3.91E-02 |
| GSE1460 DP VS CD4 THYMOCYTE DN | 192 | 23 | 2.77E-04 | 3.97E-02 |
| GSE40666 WT VS STAT1 KO CD8 TCELL UP | 190 | 25 | 2.79E-04 | 3.99E-02 |

| GSEA pathway (v.7.2) | N | DM | P-value | FDR |
|---|---|---|---|---|
| HP ABNORMALITY OF THE LYMPH NODES | 174 | 21 | 2.96E-04 | 4.21E-02 |
| GSE21033 1H VS 24H POLYIC STIM DC UP | 159 | 22 | 2.98E-04 | 4.21E-02 |
| GSE17974 2H VS 72H UNTREATED IN VITRO CD4 TCELL DN | 188 | 19 | 3.02E-04 | 4.25E-02 |
| SIG BCR SIGNALING PATHWAY | 45 | 11 | 3.12E-04 | 4.27E-02 |
| GSE11057 PBMC VS MEM CD4 TCELL DN | 183 | 22 | 3.08E-04 | 4.27E-02 |
| GSE26495 NAIVE VS PD1HIGH CD8 TCELL DN | 193 | 25 | 3.12E-04 | 4.27E-02 |
| GSE21670 STAT3 KO VS WT CD4 TCELL TGFB IL6 TREATED DN | 192 | 24 | 3.06E-04 | 4.27E-02 |
| GSE20727 CTRL VS H2O2 TREATED DC UP | 192 | 24 | 3.11E-04 | 4.27E-02 |
| GO RESPONSE TO MUSCLE INACTIVITY | 11 | 5 | 3.07E-04 | 4.27E-02 |
| GSE22886 CD8 TCELL VS BCELL NAIVE UP | 193 | 23 | 3.17E-04 | 4.31E-02 |
| GSE22886 NAIVE TCELL VS MONOCYTE DN | 193 | 23 | 3.20E-04 | 4.31E-02 |
| GSE22432 CDC VS COMMON DC PROGENITOR UP | 188 | 24 | 3.20E-04 | 4.31E-02 |
| GO NEGATIVE REGULATION OF RESPONSE TO STIMULUS | 1670 | 111 | 3.19E-04 | 4.31E-02 |
| GSE17974 0H VS 4H IN VITRO ACT CD4 TCELL UP | 174 | 22 | 3.23E-04 | 4.32E-02 |
| GSE2935 UV INACTIVATED VS LIVE SENDAI VIRUS INF MACROPHAGE UP | 164 | 23 | 3.24E-04 | 4.32E-02 |
| ZNF669 TARGET GENES | 112 | 14 | 3.25E-04 | 4.32E-02 |
| GO CELLULAR RESPONSE TO HORMONE STIMULUS | 611 | 52 | 3.34E-04 | 4.42E-02 |
| GSE28726 NAIVE VS ACTIVATED CD4 TCELL UP | 191 | 22 | 3.38E-04 | 4.46E-02 |
| TONKS TARGETS OF RUNX1 RUNX1T1 FUSION HSC DN | 181 | 22 | 3.51E-04 | 4.51E-02 |
| MIR12133 | 365 | 40 | 3.48E-04 | 4.51E-02 |
| GSE1460 INTRATHYMIC T PROGENITOR VS NAIVE CD4 TCELL CORD BLOOD DN | 190 | 22 | 3.46E-04 | 4.51E-02 |
| GSE26495 NAIVE VS PD1LOW CD8 TCELL DN | 193 | 26 | 3.51E-04 | 4.51E-02 |
| GO NEGATIVE REGULATION OF HEMOPOIESIS | 150 | 17 | 3.46E-04 | 4.51E-02 |
| WP MICRORNAS IN CARDIOMYOCYTE HYPERTROPHY | 99 | 15 | 3.49E-04 | 4.51E-02 |
| GOLDRATH ANTIGEN RESPONSE | 338 | 28 | 3.60E-04 | 4.54E-02 |
| CAGCTTT MIR320 | 231 | 27 | 3.56E-04 | 4.54E-02 |
| GSE29618 BCELL VS MDC UP | 176 | 22 | 3.61E-04 | 4.54E-02 |
| GSE7218 IGM VS IGG SIGNAL THGOUGH ANTIGEN BCELL DN | 169 | 17 | 3.61E-04 | 4.54E-02 |
| GSE7219 WT VS NIK NFKB2 KO DC DN | 160 | 21 | 3.56E-04 | 4.54E-02 |
| WP TCELL RECEPTOR AND COSTIMULATORY SIGNALING | 29 | 8 | 3.62E-04 | 4.54E-02 |
| FAN EMBRYONIC CTX BIG GROUPS BRAIN IMMUNE | 144 | 18 | 3.64E-04 | 4.54E-02 |
| GSE22886 DAY0 VS DAY7 MONOCYTE IN CULTURE UP | 188 | 22 | 3.66E-04 | 4.54E-02 |
| GSE7460 FOXP3 MUT VS WT ACT WITH TGFB TCONV DN | 193 | 25 | 3.67E-04 | 4.54E-02 |
| GO ALPHA BETA T CELL ACTIVATION INVOLVED IN IMMUNE RESPONSE | 61 | 10 | 3.86E-04 | 4.77E-02 |
| GSE7460 TCONV VS TREG THYMUS DN | 194 | 21 | 3.92E-04 | 4.81E-02 |
| GSE22601 IMMATURE CD4 SINGLE POSITIVE VS DOUBLE POSITIVE THYMOCYTE DN | 196 | 23 | 3.97E-04 | 4.82E-02 |
| GO NEGATIVE REGULATION OF PROTEIN MODIFICATION PROCESS | 611 | 50 | 3.96E-04 | 4.82E-02 |
| GO LEUKOCYTE MIGRATION | 436 | 36 | 3.97E-04 | 4.82E-02 |
| GSE24142 EARLY THYMIC PROGENITOR VS DN2 THYMOCYTE UP | 195 | 24 | 4.00E-04 | 4.85E-02 |
| PID ANGIOPOIETIN RECEPTOR PATHWAY | 47 | 10 | 4.06E-04 | 4.87E-02 |
| MIR1271 5P | 368 | 40 | 4.09E-04 | 4.87E-02 |
| GSE360 LOW DOSE B MALAYI VS M TUBERCULOSIS MAC UP | 189 | 23 | 4.06E-04 | 4.87E-02 |
| GSE7852 TREG VS TCONV LN UP | 188 | 24 | 4.09E-04 | 4.87E-02 |
| GSE6259 FLT3L INDUCED 33D1 POS DC VS BCELL DN | 188 | 23 | 4.10E-04 | 4.87E-02 |
| MIR3166 | 110 | 17 | 4.18E-04 | 4.88E-02 |

| GSEA pathway (v.7.2) | N | DM | P-value | FDR |
|---|---|---|---|---|
| LYF1 01 | 255 | 28 | 4.20E-04 | 4.88E-02 |
| GSE13411 IGM MEMORY BCELL VS PLASMA CELL UP | 184 | 22 | 4.25E-04 | 4.88E-02 |
| GSE31082 DP VS CD4 SP THYMOCYTE DN | 190 | 22 | 4.14E-04 | 4.88E-02 |
| GSE360 CTRL VS T GONDII MAC UP | 191 | 21 | 4.24E-04 | 4.88E-02 |
| GSE3039 ALPHABETA CD8 TCELL VS B2 BCELL UP | 192 | 24 | 4.22E-04 | 4.88E-02 |
| GSE21774 CD62L POS CD56 DIM VS CD62L NEG CD56 DIM NK CELL UP | 190 | 24 | 4.24E-04 | 4.88E-02 |
| GSE28726 ACT CD4 TCELL VS ACT NKTCELL DN | 191 | 22 | 4.18E-04 | 4.88E-02 |
| GO POSITIVE REGULATION OF ALPHA BETA T CELL DIFFERENTIATION | 44 | 9 | 4.23E-04 | 4.88E-02 |
| HOEBEKE LYMPHOID STEM CELL UP | 94 | 17 | 4.27E-04 | 4.88E-02 |
| ODONNELL TARGETS OF MYC AND TFRC UP | 74 | 11 | 4.35E-04 | 4.95E-02 |
| GO POSITIVE REGULATION OF RESPONSE TO EXTERNAL STIMULUS | 505 | 38 | 4.36E-04 | 4.95E-02 |
| GSE11057 CD4 CENT MEM VS PBMC UP | 192 | 23 | 4.41E-04 | 4.95E-02 |
| GSE1448 ANTI VALPHA2 VS VBETA5 DP THYMOCYTE DN | 188 | 21 | 4.40E-04 | 4.95E-02 |
| GO POSITIVE REGULATION OF LYMPHOCYTE DIFFERENTIATION | 99 | 14 | 4.39E-04 | 4.95E-02 |

Notes: N: is the number of genes contained in the curated pathway, DM is the number of differentially methylated genes corrected for probes assigned to multiple genes or transcripts of the same gene, P-value is the p-value for the hypergeometric test of overrepresentation, and FDR is the Q-value false discovery rate to compare for multiple comparisons.

Supplementary Table 8. Baseline characteristics of the samples included in the application datasets from GEO and ArrayExpress

| Dataset (Platform) | N | mean Age (sd) | n male (%) | Sample | Data Source |
|---|---|---|---|---|---|
| **DISEASES** | | | | | |
| **Multiple Sclerosis (450k)** | | | | | |
| Case | 13 | NA | NA | whole blood | GSE88824 |
| Control | 14 | NA | NA | whole blood | GSE88824 |
| **Rheumatoid arthritis (450k)** | | | | | |
| Case | 354 | 51.15 (12.05) | 101 (28.5) | peripheral blood leukocytes | GSE42861 |
| Control | 355 | 52.76 (11.48) | 96 (28.7) | peripheral blood leukocytes | GSE42861 |
| **Breast Cancer Treatment (EPIC)** | | | | | |
| Radiation-therapy | 74 | 57.38 (9.24) | 0 (0) | peripheral blood | GSE140038 |
| Radiation-therapy and chemotherapy | 70 | 56.34 (11.15) | 0 (0) | peripheral blood | GSE140038 |
| **COVID-19 (EPIC)** | | | | | |
| Healthy | 6 | 55.8 (6.15) | NA | peripheral blood | GSE161778 |
| No remission | 2 (3 samples | 64.67 (13.28) | NA | peripheral blood | GSE161778 |
| Remission | 4 (15 samples) | 65.73 (12.79) | NA | peripheral blood | GSE161778 |
| **SUBJECT-TO-SUBJECT VARIATION** | | | | | |
| **Twin (450k)** | | | | | |
| Monozygotic twins | 852 | NA | 438 (51.4) | whole blood | GSE105018 |
| Dizygotic twins | 612 | NA | 312 (50.1) | whole blood | GSE105018 |
| **Aging (450k+EPIC)** | | | | | |
| Newborn | 141 | 0.00 (0.00) | 62 (44.0) | cord blood | E-MTAB-7069, GSE85042, GSE103189, GSE104778 |
| 0-5 | 71 | 2.73 (1.87) | 6 (8.5) | peripheral blood | E-MTAB-7069, GSE62219 |
| 5-18 | 95 | 15.71 (2.33) | 47 (49.5) | peripheral blood | E-MTAB-7069, GSE87571 |
| 18-65 | 1049 | 49.93 (12.43) | 317 (30.2) | peripheral blood | E-MTAB-7309, GSE87571, GSE121633, |
| >65 | 1148 | 75.75 (7.07) | 408 (35.5) | peripheral blood | E-MTAB-7309, GSE87571, GSE121633, |
| **Total** | 4872 | | | | |

Supplementary Fig. 8. Predicted immune cell proportions in whole blood samples between multiple sclerosis cases (n=13) and normal controls (n=14) (450k)

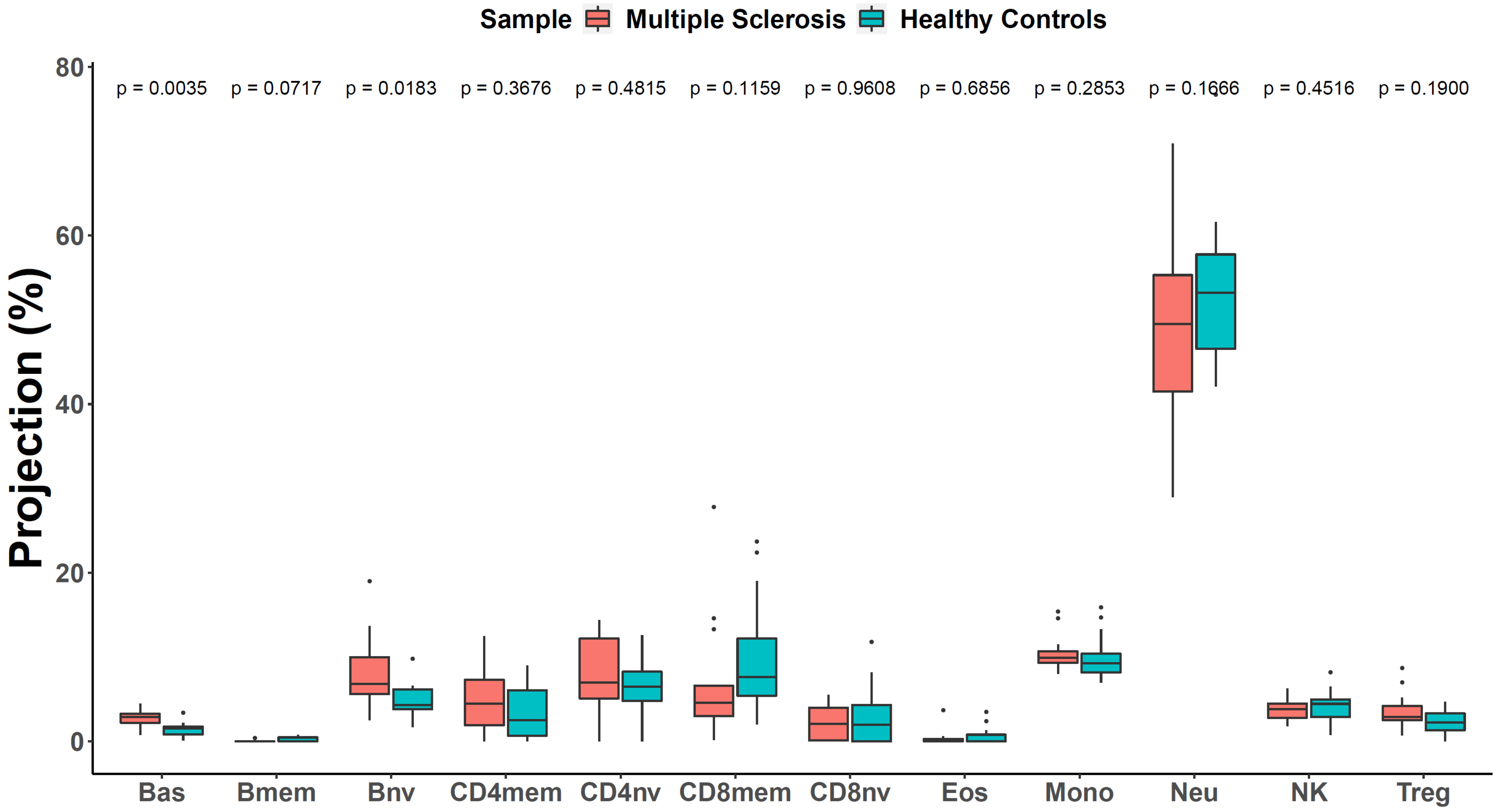

Supplementary Fig. 9. Predicted immune cell proportions in peripheral blood leukocyte samples between rheumatoid arthritis cases (n=354) and normal controls (n=355) (450k)

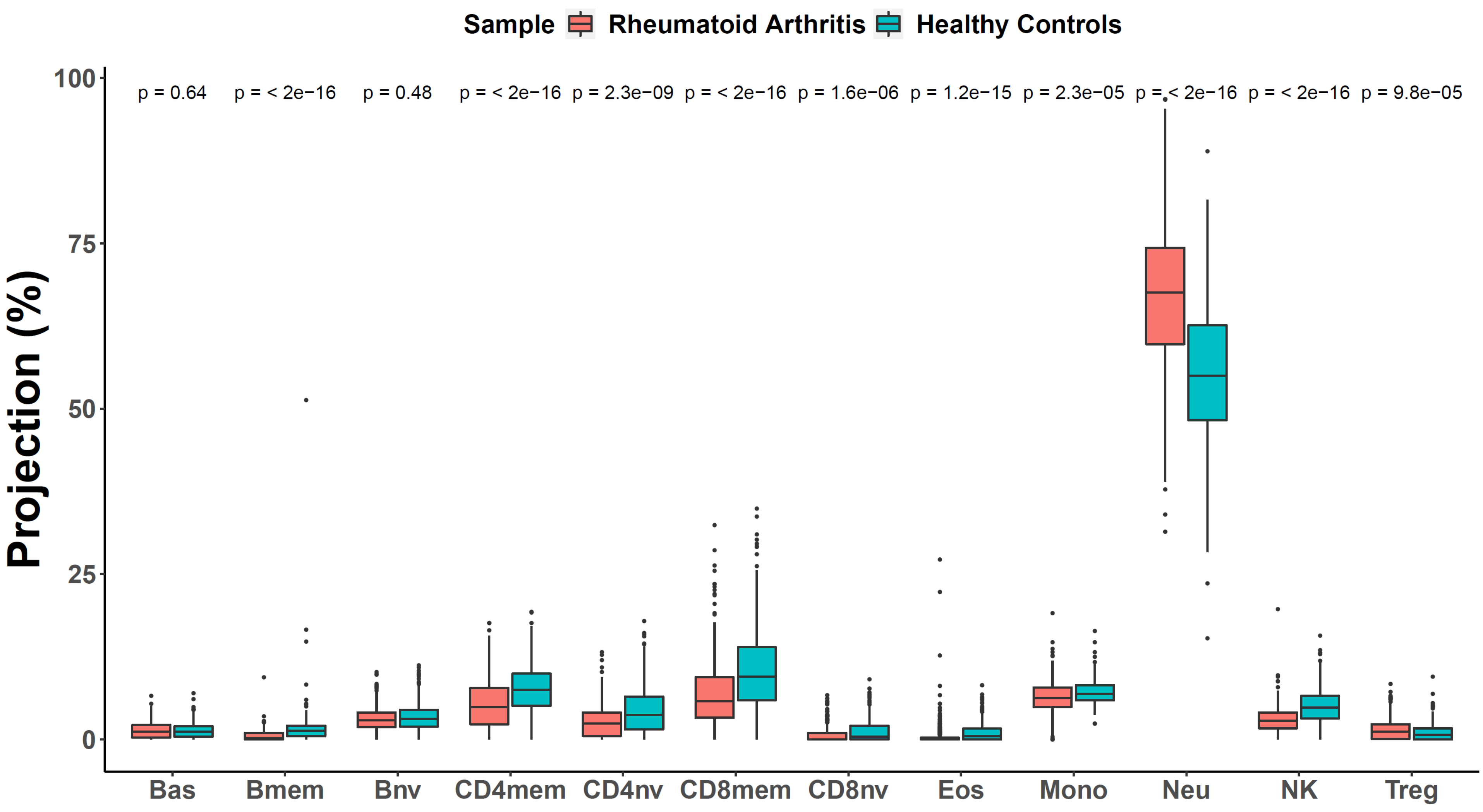

Supplementary Fig. 10. Predicted immune cell proportions in peripheral blood samples from early breast cancer patients before and after receiving radiation therapy only (n=74) and radiation therapy plus chemotherapy (n=70) (EPIC)

Panel A

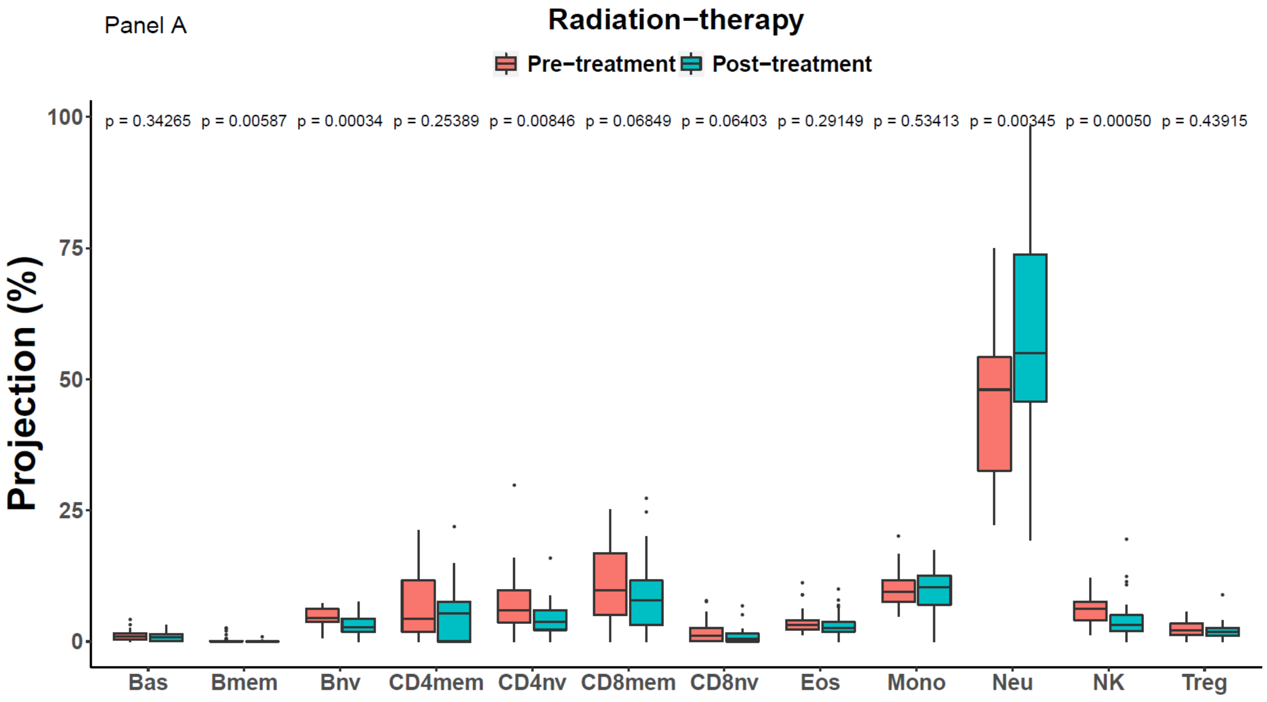

Panel B

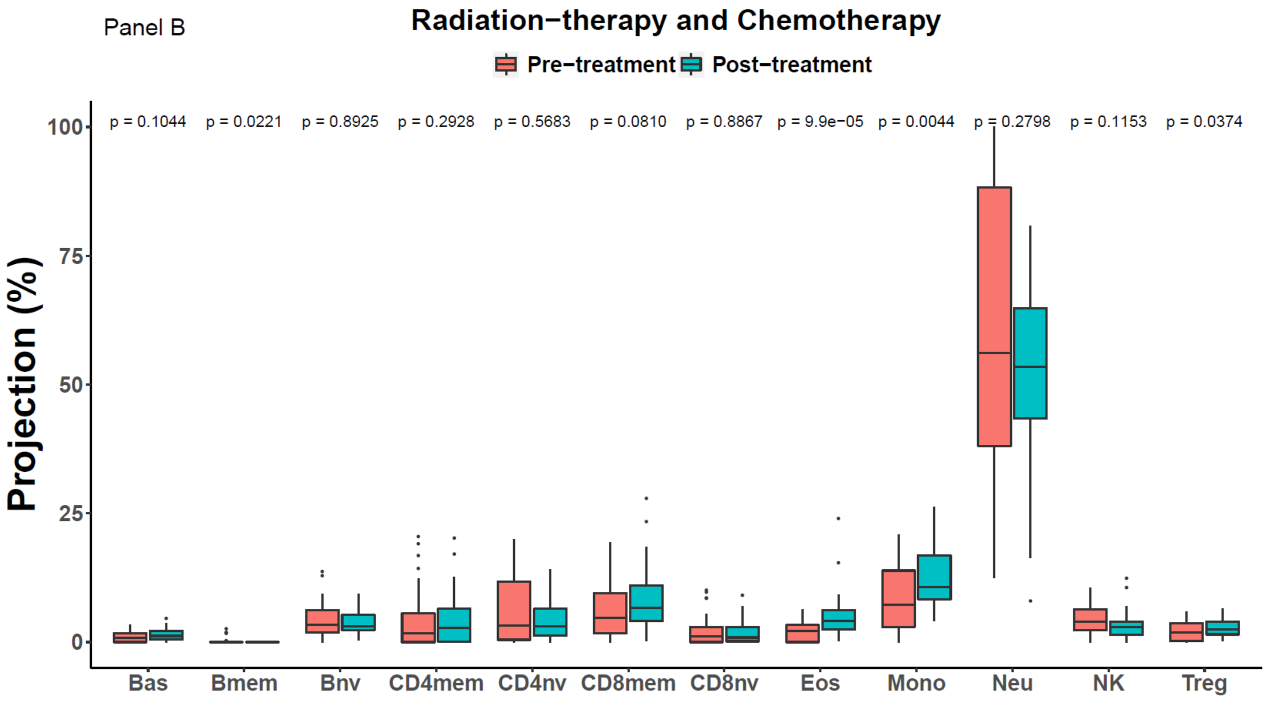

Supplementary Fig. 11. Changes in estimated immune cell proportions between subjects with COVID-19 infection with and without remission compared to healthy subjects (EPIC)

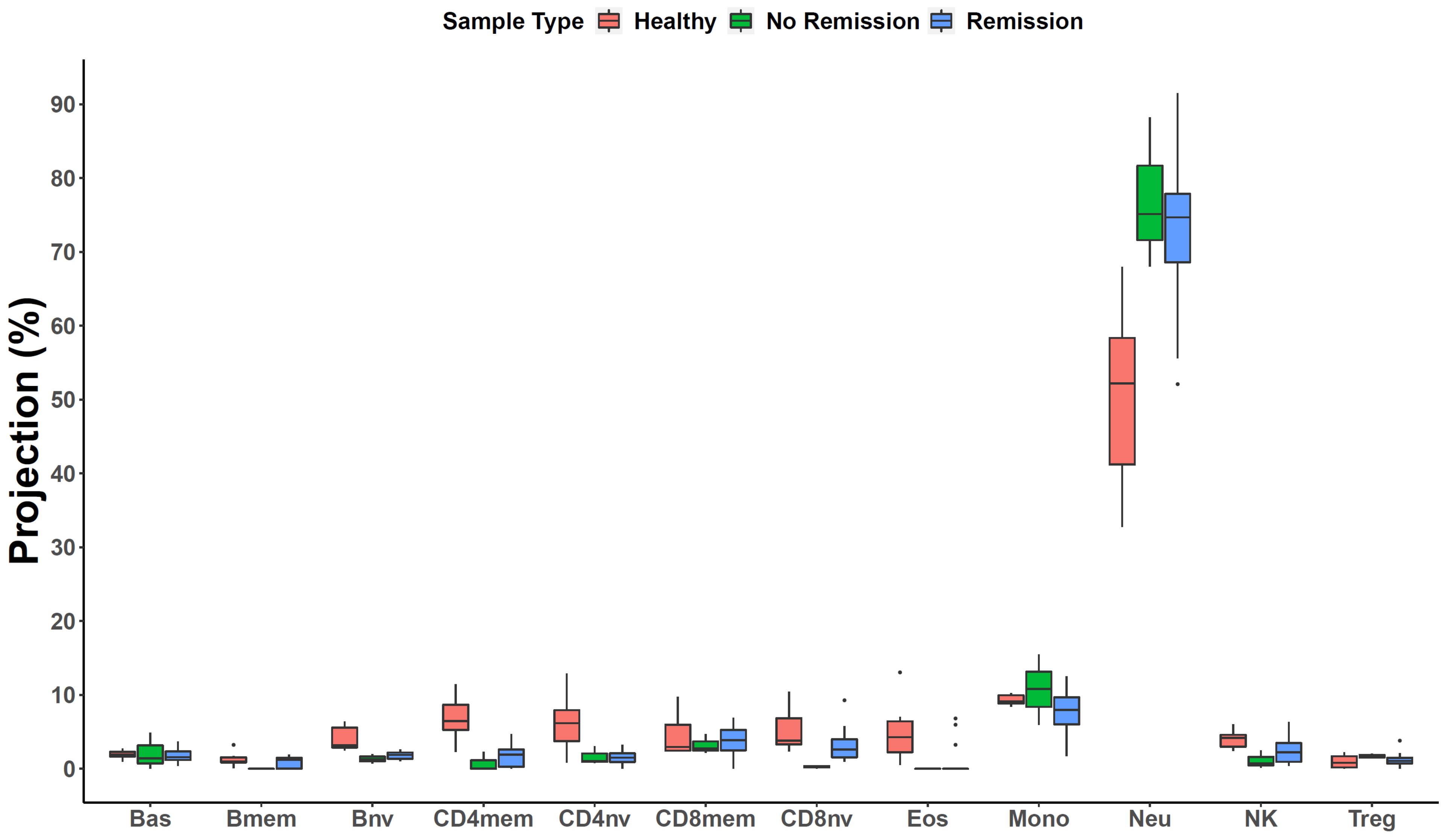

Supplementary Fig. 12. Differences of predicted immune cell proportions in whole blood samples between pairs of twins in monozygotic twins (n=852) and dizygotic twins (n=612) (450k)

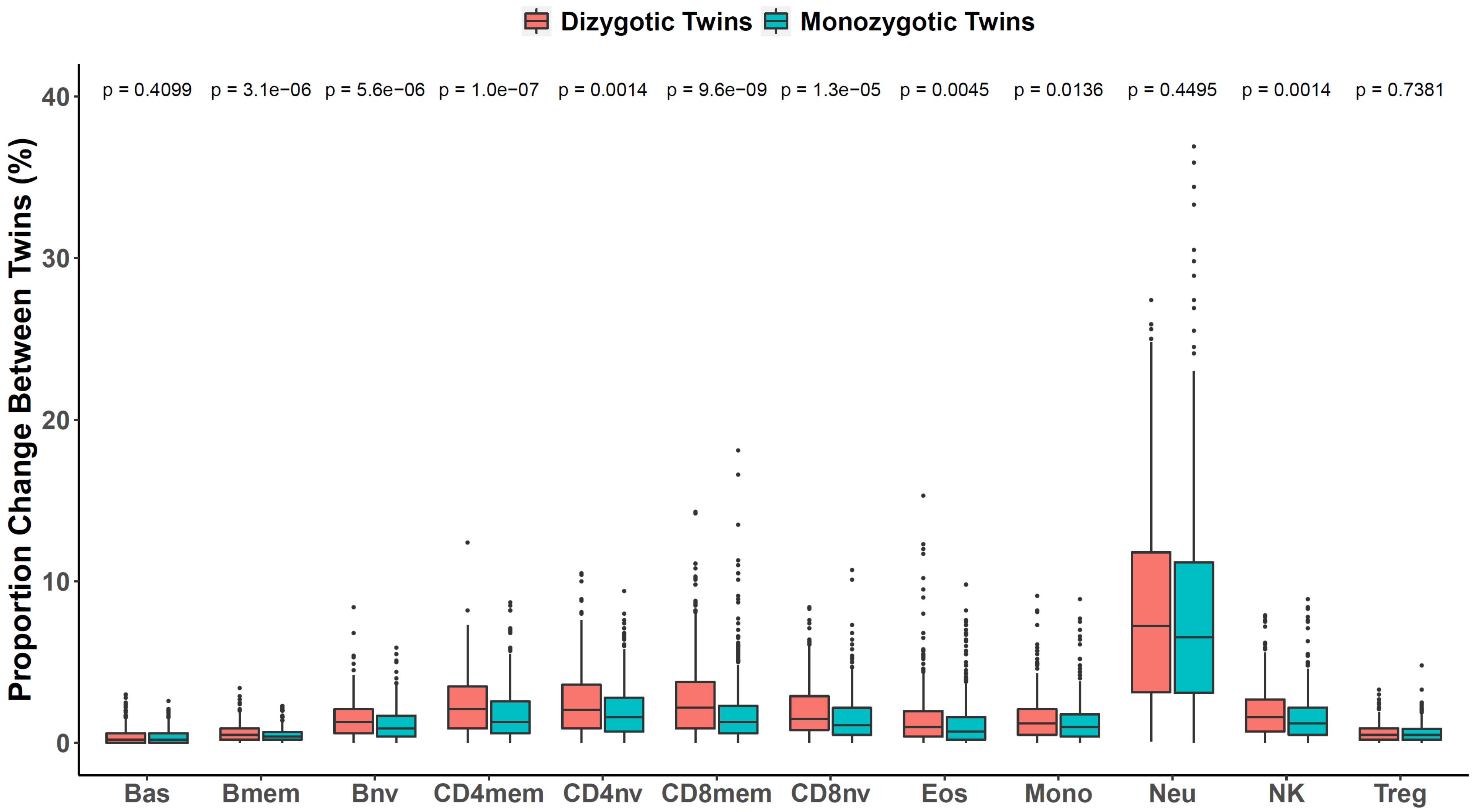

Supplementary Fig. 13. Trajectories of several cell subpopulation ratios across different ages using publicly available datasets (n=2504)

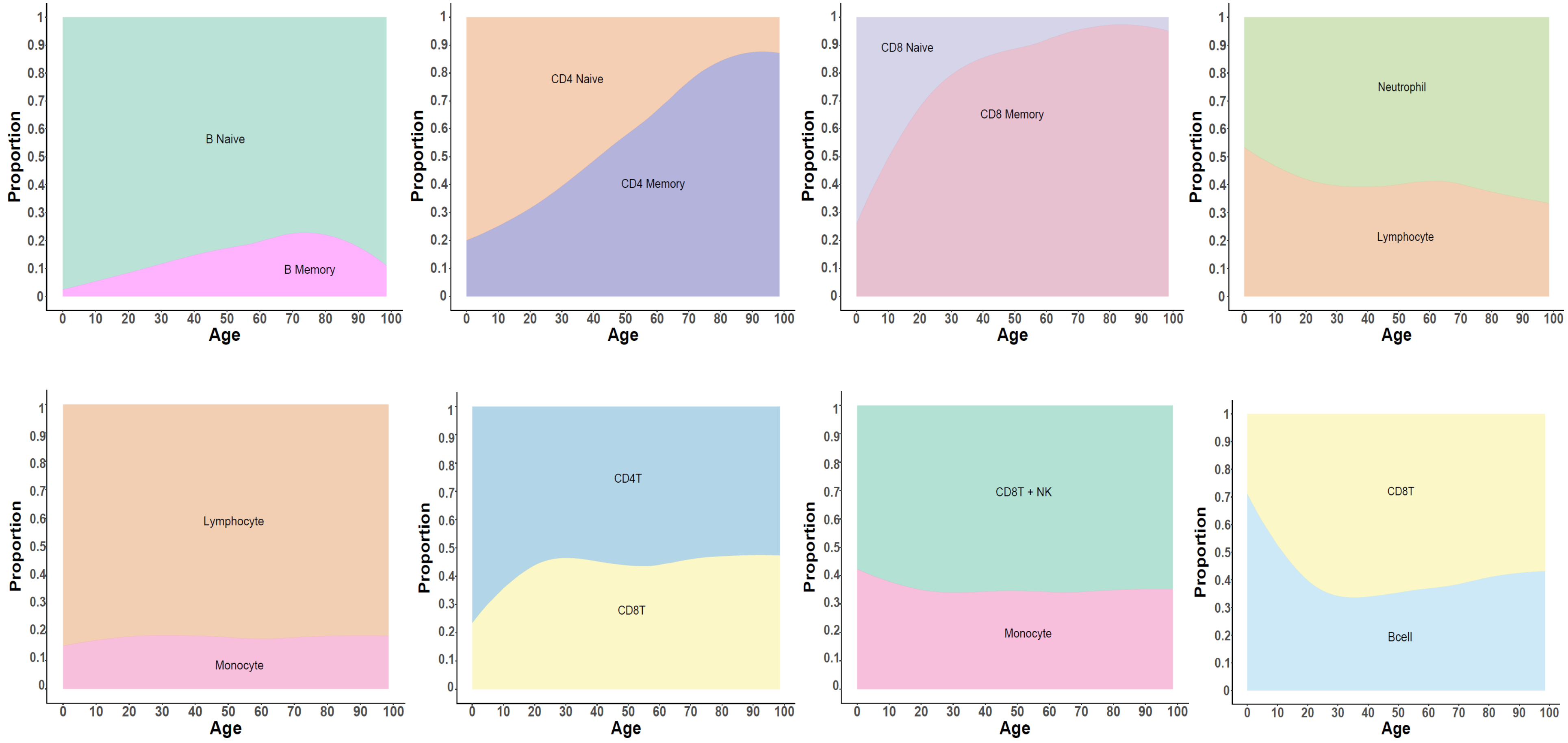

Notes: The proportion of the estimated cells were averaged and smoothed to represent the proportion for the compartment and the ratio between the subpopulations for specific ages. The subjects represent “healthy controls”, however, it is unknown if specific subjects had any comorbidity at the time of the sample array (For granular representation fo the data see the Supplementary Figure S11).

Supplementary Fig. 14. Changes on predicted immune cell proportions with aging using samples from ages zero to greater than 90 years (n=2504) (450k and EPIC)

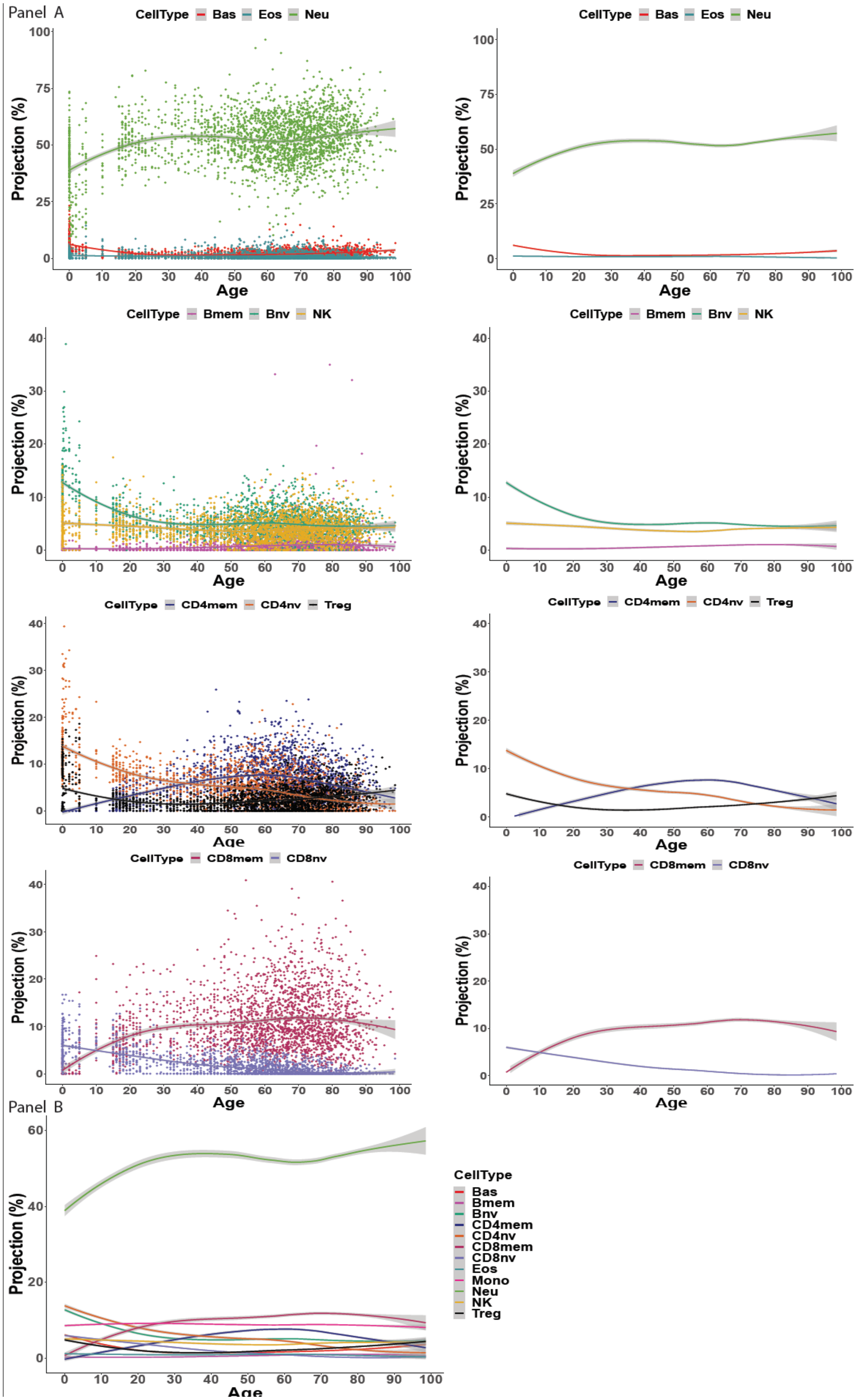

Supplementary Fig. 15. Longitudinal changes of predicted immune cell proportions within 5 years after birth in human blood leukocytes of 10 healthy girls

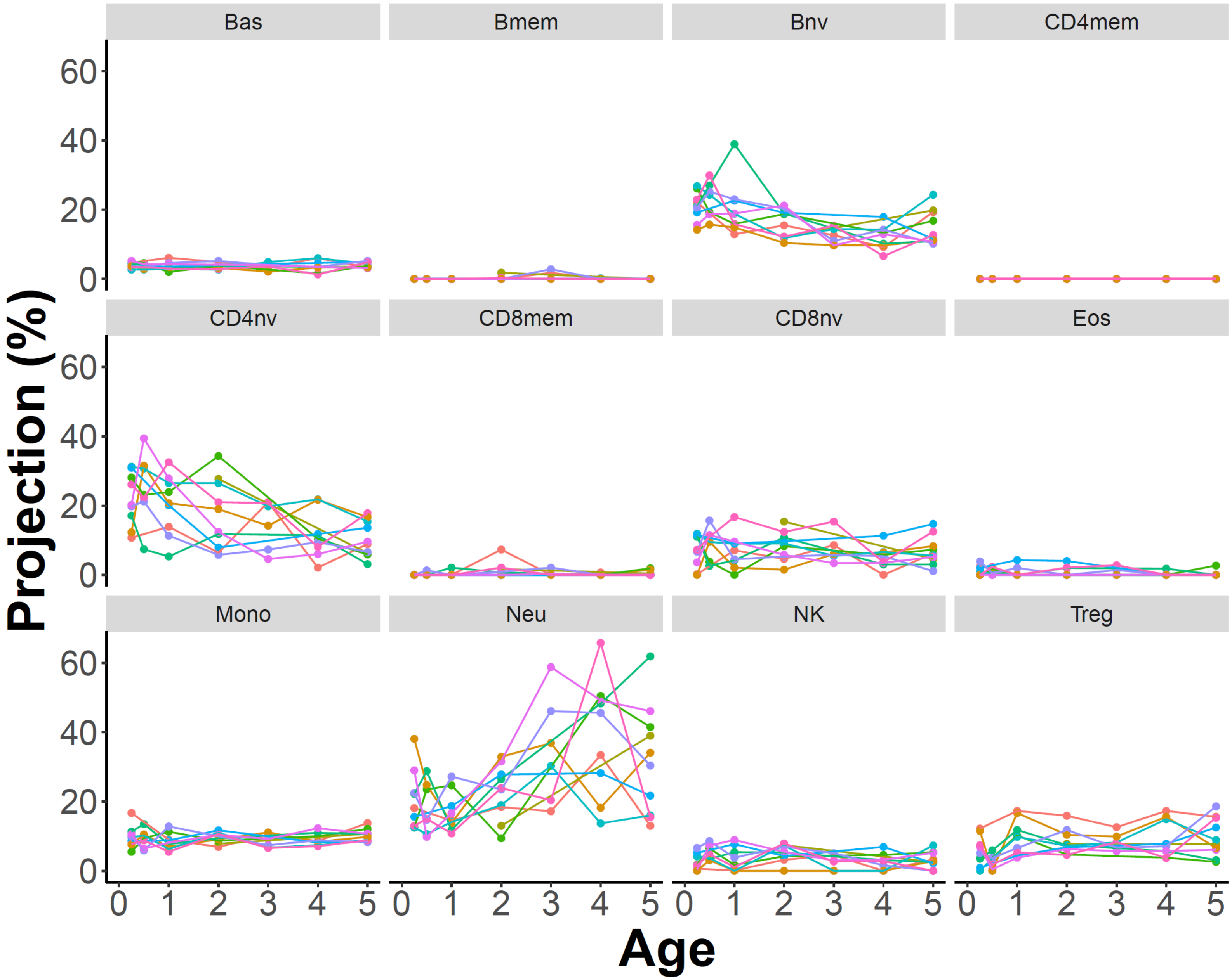
